# Structural Premise of Selective Deubiquitinase USP30 Inhibition by Small-Molecule Benzosulfonamides

**DOI:** 10.1101/2022.09.13.507798

**Authors:** Darragh P O’Brien, Hannah BL Jones, Franziska Guenther, Emma J Murphy, Katherine S England, Malcolm Anderson, Paul Brennan, John B Davis, Adán Pinto-Fernández, Andrew P Turnbull, Benedikt M Kessler

## Abstract

Dampening functional levels of the mitochondrial deubiquitylating enzyme USP30 has been suggested as an effective therapeutic strategy against neurodegenerative disorders such as Parkinson’s Disease. USP30 inhibition may counteract the deleterious effects of impaired turnover of damaged mitochondria which is inherent to both familial and sporadic forms of the disease. Small-molecule inhibitors targeting USP30 are currently in development, but little is known about their precise nature of binding to the protein. We have integrated biochemical and structural approaches to gain novel mechanistic insights into USP30 inhibition by a small-molecule benzosulfonamide containing compound, **39**. Activity-based protein profiling (ABPP) mass spectrometry confirmed target engagement, the high selectivity, and potency of **39** for USP30 against 49 other deubiquitylating enzymes in a neuroblastoma cell line. *In vitro* characterization of **39** enzyme kinetics infers slow and tight binding behavior, which is comparable with features of covalent modification of USP30. Finally, we blended hydrogen-deuterium exchange mass spectrometry and computational docking to elucidate the molecular architecture and geometry of USP30 complex formation with **39**, identifying structural rearrangements at the cleft of the USP30 thumb and palm subdomains. These studies suggest that **39** binds to the thumb-palm cleft that guides the ubiquitin C-terminus into the active site, thereby preventing ubiquitin binding and isopeptide bond cleavage, and confirming its importance in the inhibitory process. Our data will pave the way for the design and development of next-generation inhibitors targeting USP30 and associated deubiquitinylases.

## INTRODUCTION

Ubiquitination is essential to protein quality control, homeostasis and lifespan.^1^ Damaged proteins are flagged for removal from cells with the covalent addition of ubiquitin (Ub), a small, highly conserved 76-amino acid protein that is widely expressed across eukaryotic cell types.^2, 3^ This molecular “kiss of death” proceeds through the coordinated action of E1, E2, and E3 ligase enzymes, with targeted protein substrates conjugated by way of an isopeptide bond to either a single Ub molecule (mono-ubiquitination), or several repeating units (poly-ubiquitination).^4^ Besides the Ub-proteasome system (UPS) subfamily, proteins selected for degradation can also be trafficked through the autophagy-lysosome pathway. In damaged mitochondria, such autophagic clearance of impaired proteins occurs through a highly selective and dedicated mechanism termed “mitophagy”. Several key players work tirelessly in the dysregulated mitochondrion to maintain overall cell integrity and survival, including the mitochondrial outer membrane (MOM)-associated Ub serine/threonine kinase PINK1 and the cytoplasmic E3 ligase Parkin.^5^ PINK1 phosphorylates Ub species on damaged proteins accumulating on the MOM, flagging them for elimination. This in turn recruits and activates endogenous Parkin (also by phosphorylation), initiating a hyper-ubiquitination cascade that gives the green light for mitophagy to proceed in a specialized autophagosome structure.^6^ Deubiquitinating enzymes (DUBs) counteract the actions of E3 ligases by removing Ub modifications, and several of these Ub-specific proteases (USPs) have been shown to oppose Parkin activity.^7, 8^ Of these, USP30 is the predominant active DUB to be directly implicated in mitophagy to-date, primarily due to its localization on the MOM, whilst also being linked to pexophagy and oxygen metabolism due to its widespread expression on peroxisomes.^9^ Interestingly, Parkin and USP30 both share an unusual preference for Lys6-linked Ub chains, the mitophagic importance of which has yet to be conclusively deciphered.^10, 11^

Impaired mitophagy and oxidative stress have adverse roles in neurodegeneration, with both being linked to familial and sporadic forms of Parkinson’s Disease (PD)^12, 13^; PINK1/Parkin-mediated mitophagy limits the build-up of toxic mitochondrial species in PD and loss-of-function mutations occurring in the PINK1 and PRKN genes result in the progressive depletion of dopaminergic neurons of the basal ganglia and a rare hereditary form of juvenile Parkinsonism.^14, 15^ As USP30 antagonizes mitophagy through ubiquitination, its inhibition has been proposed as a novel therapeutic strategy to enhance mitochondrial turnover and clear damaged mitochondrial proteins, providing a much-needed strategy to improve outcomes in PD and other neurodegenerative disorders. Several small-molecule inhibitors targeting USP30 are in the pipeline, including phenylalanine derivatives, *N*-cyano pyrrolidines and natural products.^16–18^ Perhaps those with the greatest potential, however, are a family of benzosulfonamides, most notably, compound **39**, which has been shown to boost mitophagy in dopaminergic neurons of PD patients by down-regulating USP30.^19, 20^ Little is currently known, however, regarding the intricacies of its non-covalent attachment to USP30 itself, and the structural basis of its inhibitory action. X-ray crystallography has recently provided structures of both human and zebrafish USP30, but these are solely in the context of attachment to Lys6-linked di-Ub moieties.^10, 11^ Hydrogen/Deuterium eXchange Mass Spectrometry (HDX-MS) is a complementary biophysical tool that can provide unique insights into protein structure, stability, dynamics and function.^21^ In direct contrast to the static snapshot provided by the crystal structure, HDX-MS monitors the conformational dynamics of a system *in solution*, enabling the analysis of “proteins in motion”. The technique has shown particular utility for the rapid and reliable identification of small-molecule binding pockets on proteins.^22^ Whilst lacking the resolution of a crystal structure, the highly informative data obtained from HDX-MS can be combined with orthogonal structural, computational, biochemical and/or biophysical techniques to define structure activity relationships (SAR) and to direct drug discovery campaigns.

As such, we have integrated several biophysical and structural approaches to help clarify the molecular and structural interplay of **39** binding to USP30. The endogenous cellular selectivity of **39** for USP30 inhibition was confirmed using activity-based protein profiling mass spectrometry (ABPP-MS) against a panel of 49 other endogenous DUBs in neuronal SH-SY5Y cells. Bio-layer interferometry showed that **39** binds to USP30 in a slow and tight manner, which intriguingly, despite being a non-covalent binder, is consistent with the profile of covalent inhibition. Finally, we combined HDX-MS and molecular docking simulations to elucidate the conformational dynamics and spatial preferences of **39** binding, identifying key residues in the inhibitory process itself. **39** induces conformational and structural rearrangements at the cleft of the USP30 thumb and palm subdomains, in a region encompassing its catalytic residues and the site of Ub binding. We postulate that these phenomena underlie the mechanism of inhibition of **39**. Dampening USP30 pharmacologically may represent a tractable treatment for PD and other mitophagy-related disorders. As no previous attempts to investigate the molecular basis of compound **39** binding to USP30 have been reported, our data will be instrumental in the development of rational next-generation inhibitors against USP30 and related DUBs.

## RESULTS AND DISCUSSION

### Compound 39 is highly potent and selective for neuronal USP30

The efficacy and selectivity of compound **39** across a panel of cysteine active DUBs was initially screened in SH-SY5Y neuroblastoma cell lysates by ABPP (**Figure 1a**). SH-SY5Y cell extracts were treated with a range of inhibitor concentrations from 0.1 to 25 μM, followed by incubation with a HA-tagged Ub-based probe with a propargylamine warhead (HA-Ub-PA). DUB probe complexes were immunoprecipitated by way of their HA tag and quantified using label-free quantitation (LFQ) LC-MS/MS. We implemented a data independent acquisition (DIA) MS regime to maximize the depth and reproducibility of the DUB profiling assay.^23, 24^ The concentration-dependent competition between compound **39** and HA-Ub-PA for binding to USP30 confirmed target engagement and potency of the inhibitor in a cellular matrix, with an IC_50_ value in the nanomolar range (**Figure 1b**). Moreover, **39** was found to be highly selective for USP30 as it had no significant activity against any of the other 49 endogenous DUBs detected in the experiment (**Figure 1b**). The main cysteine-reactive DUB enzyme families were all represented, with proteins containing USP, ovarian tumor protease (OTU), Ub C-terminal hydrolase (UCH), and Josephin domains quantified.^25, 26^ The absence of **39** concentration-dependent inhibition for all other identified cysteine active DUBs demonstrates the highly selective nature of the inhibitor. This selectivity is in line with previously published USP30 inhibitor selectivity data from both a recombinant DUB activity panel^20^ and an ABPP-MS experiment on a smaller panel of endogenous DUBs identified from mouse brain tissue.^27^

**Figure 1.**
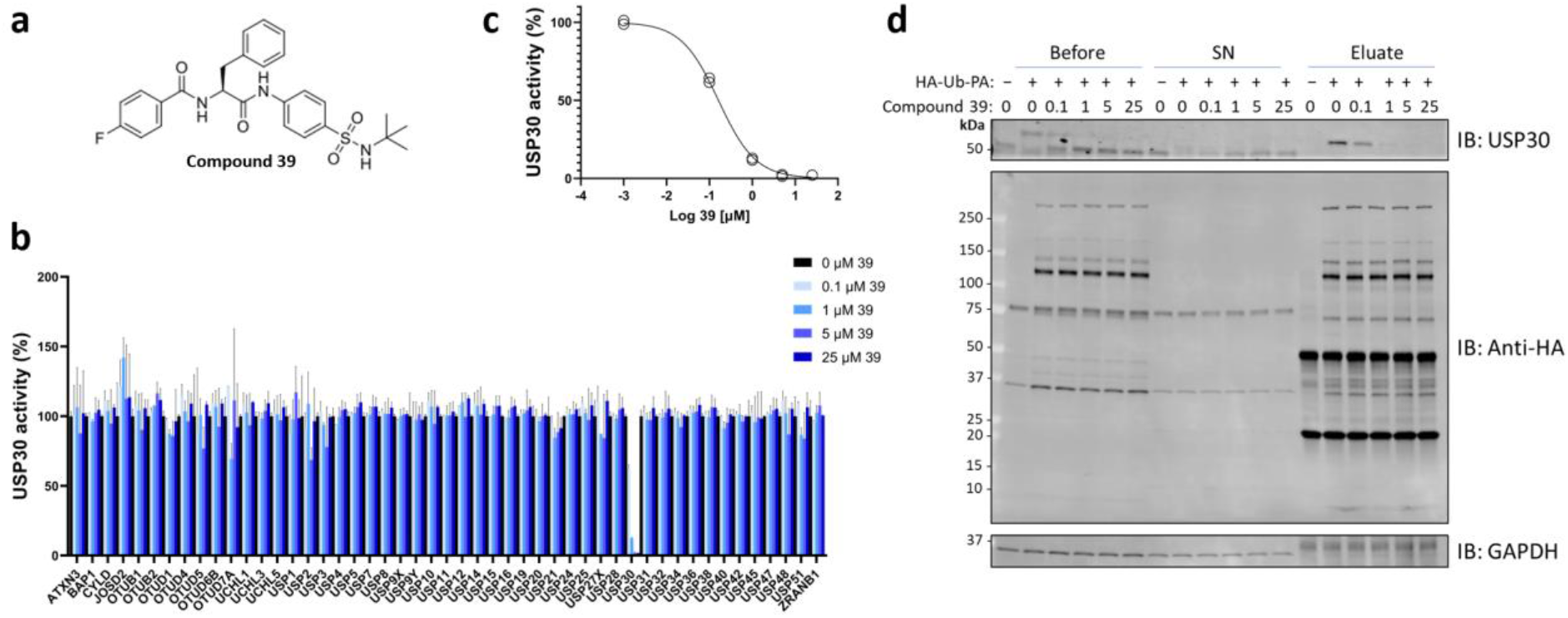
Compound 39 is highly potent and selective for USP30 inhibition in a cellular context. (a) Chemical structure of **39**.^27^ (b) Compound **39** is a highly selective USP30 inhibitor. (c) **39** acts in a concentration dependent manner (down to 0.1 μM levels), which is in concordance with previously published data for both mouse brain and with a recombinant DUBprofiler™ (Ubiquigent) panel. (d) SHSY5Y cells were incubated with **39** for 1 h at 37°C. The HA-Ub-PA was then incubated with **39**-treated lysates for 45 min at 37°C at a protein ratio of 1:200 (w/w) and immunoblotted as shown. Engagement of USP30 with the probe was confirmed using anti-USP30 and anti-HA antibodies.

We have recently reported that **39** can be displaced by HA-Ub-PA over long incubations.^27^ Accordingly, a small amount of displacement during the 45 min HA-Ub-PA incubation was anticipated. It is therefore expected that the IC_50_ value of 0.16 μM that we obtained from our ABPP-MS assay is likely higher than the absolute IC_50_ inhibition concentration (**Figure 1c** and **d**).

### Compound 39 binds USP30 in a slow and tight manner

Once it was established that **39** down-regulates endogenous USP30 activity in a highly selective fashion, we sought to rigorously profile its inhibitory properties using a recombinant version of the protein. Synthetic full-length USP30 is very unstable and difficult to solubilize.^10^ To circumvent this, we used a previously described truncated version of USP30 in our enzyme (and HDX-MS) assays, which readily went into solution and was determined to be stable over the time course of our experiments (**Figure S1**). To assess enzyme deubiquitinating efficiency, the purified USP30 construct was incubated with a fluorogenic Ub-rhodamine substrate in both the presence and absence of compound **39**. This resulted in a calculated IC_50_ value for **39** of ~2 nM *in vitro*, which was in-line (albeit 10-fold lower) with previous estimations (**Figure 2a**).^19^ Although both measurements remain in the nM range, the 2 nM IC_50_ from the Ub-rhodamine assay is lower than the 162 nM IC_50_ from the ABPP. This could be attributable to differences in the endogenous and recombinant activity of USP30, non-specific inhibitor occlusion in the cellular context of the ABPP, or displacement of compound 39 in the ABPP by HA-Ub-PA. Progress curves for Ub-rhodamine cleaved by USP30 were used to calculate the rate of inhibition. The kinetic constants k_5_, k_6_ and K_i_ gave values that were indicative of slow, tight binding behavior (**Figures 2b**, **d**, **e** and **f**). The latter was visualized by a time-dependent shift of dose-response inhibition curves (**Figure 2d**), as well as by plotting IC_50_ values against time (**Figure 2e**). When considering the binding Scheme A (**Experimental Section**), the small value for k_6_ implies that it is behaving in an irreversible manner. The progress curves show typical features of an enzyme reaction in the presence of a slow binding inhibitor. Furthermore, two binding events are observed in the form of a) an initial and b) a steady-state velocity – both of which need to be considered during curve fitting and calculation of inhibitory rates (see **Equations 2** and **3** in **Supporting Information**) (**Figure 2b**).

**Figure 2.**
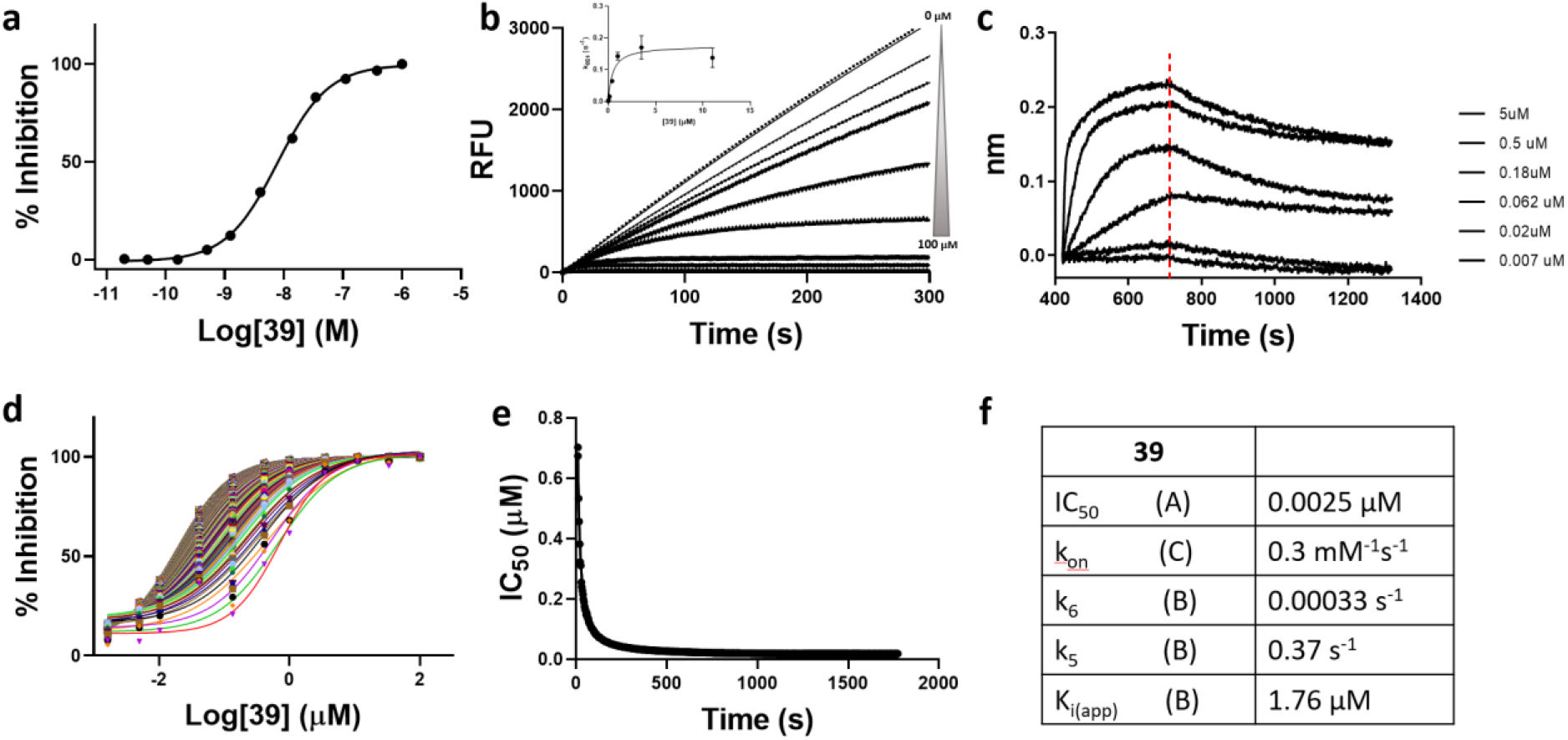
Kinetic profiling of the non-covalent USP30 inhibitor 39. Upper panel: (a) Dose dependent inhibition of USP30 by **39**. (b) Reaction progress curves recorded on the FLIPR^®^ Tetra. Traditional method for determining kinetic constants associated with a two-step slow, tight binding inhibitor. k_obs_, determined by fitting the progress curves to Eq. 2 (**Supporting Information**), is plotted vs [Compound] and fitted to Eq. 3. (**Supporting Information**) to determine K_i_, k_5_ and k_6_. (c) Bio-layer interferometry showing binding of **39** to immobilized USP30 with no detectable dissociation. Lower panel: (d) Krippendorf method^28^ (**Supporting Information**) was used as an alternative way of determining kinetic constants. Time-dependent IC_50_ curves. Each curve represents inhibition data at an individual incubation time from 3-1800 sec. (e) IC_50_ values vs. incubation time fitted to Eq. 1. (**Supporting Information**) to obtain K_i_ and k_inact_. As **39** is non-covalent compound but has a k_6_ which is essentially 0, k_inact_ in this case represents k_5_. (f) Data table of inhibition properties.

Bio-layer interferometry experiments confirmed this slow and tight binding behavior (**Figure 2c**). The compound had an association rate of 0.3 mM^−1^s^−1^ and a very slow dissociation from USP30 that was comparable with features of covalent modification (**Figure 2f**). This phenomenon is intriguing, as **39** is known to bind to USP30 (as shown by MS), by exclusively non-covalent means.

### HDX-MS kinetics identifies key residues at the compound 39 binding interface of USP30

Knowledge is currently lacking on the precise location and mechanistics of **39** binding to USP30. HDX-MS experiments were consequently designed to pinpoint the key regions of **39** binding to USP30, whilst providing novel structural insights into the solution conformation and dynamics of complex formation. HDX relies on the natural isotopic exchange of the amide backbone hydrogens of a protein with deuterium when placed in a deuterated solution.^21^ This leads to protein mass increases that are directly measurable by MS which can serve as direct probes of protein solvent accessibility and structure. Shielding of the deuterated solvent following introduction of a binding partner is indicative of a binding interface. We sought to identify such regions following **39** binding to USP30 by directly comparing the differences in HDX-MS uptake patterns of USP30 before (apo-USP30; in the presence of DMSO) and after (holo-USP30; in the presence of **39**) complex formation.

Following digestion of unlabeled USP30 with pepsin, a total of 723 peptides were generated for the protein, 133 of which were shortlisted for downstream data analysis (**Figure S2**). Selected peptides covered 96.2% of the USP30 sequence, with an average of 4.19 peptides covering each amino acid. The kinetics of deuterium uptake was analysed for all regions of USP30, which included USP domains 1-6 and the catalytic triad at Cys77, His452 and Ser477 (**Figure S3a**). From three independent replicates, the relative fractional exchange was calculated for all peptides at each of the four time points 30, 60, 600, and 3600 sec, and plotted as a function of peptide position (**Figures 3a and S3b**).

**Figure 3.**
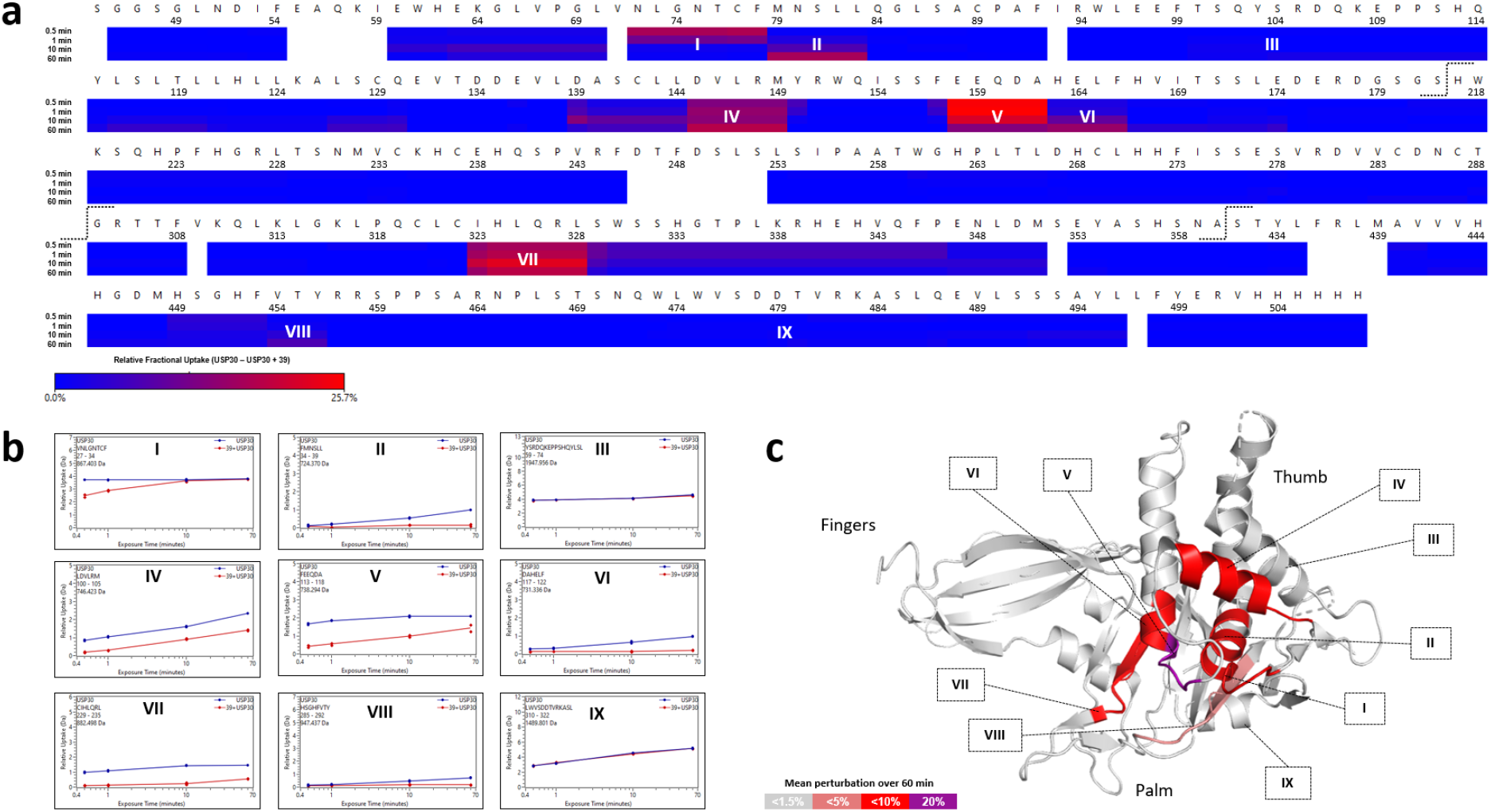
HDX-MS characterises the conformational dynamics of compound 39 binding to USP30. (a) Residue-level heat map indicating that compound **39** induces solvent protection in several regions of USP30. The plot displays the difference in relative fractional uptake between the holo- and apo-form of the protein over 1 h. Regions of which have the greatest perturbation following **39** binding are labeled Regions I-IX (b) Comparative uptake plots of Regions I-IX for apo- and holo-USP30 states (c) Integrated HDX-MS and X-Ray crystal structure of USP30 in complex with di-Ub. Regions of perturbation between apo- and holo-USP30 states HDX-MS data are colored according to magnitude of change. The data indicates that the **39** binding interface is located between the USP30 thumb and palm domains of the protein. Numbering is in accordance with the crystal structure of 5OHK. Dotted lines indicate the site of cleavage and removal of unstable, disordered sequences from the full-length USP30 protein.

Following incubation with compound **39** in conditions conducive to binary complex formation, our comparative HDX-MS data indicate that the majority of USP30 is unaffected by inhibitor binding, with no differences in deuterium uptake kinetics between apo- and holo-states (**Figure 3a**). This suggests that **39** binding is confined to smaller subsections of the protein. Indeed, several short regions of USP30 had significant shielding from the solvent in the presence of the inhibitor as compared to the DMSO control, indicative of regions involved in compound binding (**Figures 3a** and **b**). The areas of most significant perturbation included at two peptides spanning the USP30 catalytic Cys77 residue, N72-L83 (labeled Regions I and II in **Figure 3** and highlighted in representative uptake plots in **Figure 3b**), in addition to peptides mapping to D145-M149 (Region IV), E158-F166 (Regions V and VI), I323-L328 (Region VII), and finally, H449-Y456 (Region VIII), which encompasses the catalytic His452. These were in direct contrast to Regions III and IX, which gave identical HDX-MS behaviors in both the apo- and holo-form, reflecting the majority of the USP30 protein sequence (**Figure 3b**). The shielding induced by inhibitor was greatest at the 1 h timepoint, where a difference of >25% was observed in the relative fractional uptake between states. Results were confirmed by the presence of multiple overlapping peptides displaying equivalent HDX-MS activity. Interestingly, no difference in conformational dynamics was observed between states for the region covering the catalytic Ser477 (Region IX), suggesting that this site is not vital to the **39** inhibitory process, but rather, may have a greater influence in determining USP30 Ub linkage preferences.^10, 11, 29^ Nevertheless, targeting this site to improve USP30 inhibition efficacies may prove fruitful in future design regimes.

No X-ray crystal structure of apo-USP30 or USP30 in complex with **39** currently exists. We therefore mapped our solution HDX-MS data to PDB code 5OHK, which at a resolution of 2.34 Å, represents the highest resolution 3D structure of the protein currently available in the Protein Data Bank (PDB).^10^ It is worth noting that this structure corresponds to USP30 in covalent complex with Lys6-linked di-Ub, and as such, may give rise to subtle discrepancies when comparing across results. Nevertheless, due to the lack of more suitable alternatives, we felt it a worthwhile pursuit to map our apo- and holo-USP30 HDX-MS data to this 3D model. To facilitate a facile overview of the entire dataset, HDX-MS results were collapsed into a single datapoint by calculating the mean perturbation between apo- and holo-states across the four labeling time points. USP30 constitutes three subdomains designated “thumb, palm and fingers”, which is in line with related USP family members of elucidated structure.^7, 30^ Strikingly, the shielded USP30 peptides in the presence of **39** all cluster to the same spatially adjacent region of the protein, which lies at the interface of its palm and thumb subdomains (**Figure 3c**). Moreover, the regions with the greatest perturbation as highlighted above, I323-L328 (Region VII) and H449-Y456 (Region VIII), cover areas of the protein which lie opposite each other on the 3D structure. They may represent an entrance vector to the USP30 binding pocket, which is anticipated to be closer to the site of greatest perturbation at E158-F166 (Regions V and VI) and the nearby catalytic Cys77. The importance of these residues to the inhibitory process is further strengthened by their correlation with the proposed region of USP30 binding to the Ub C-terminal tail in the crystal structure.^10^

### Binding of 39 alters USP30 conformation and induces rigidification in several regions

Although likely to be in good agreement with the majority of 5OHK, the structural make-up of apo-USP30 has yet to be experimentally confirmed. As yet, no high-resolution crystal structure exists, which is of a direct consequence of the poor stability of the full-length protein itself.^10^ As stated earlier, a highly truncated USP30 construct was used in this study, which was devoid of its N-terminal mitochondrial intermembrane domain and adjacent transmembrane domain. Furthermore, several long, disordered regions were cleaved, and multiple hydrophobic residues mutated out, resulting in substantially improved protein stability and solubility. As HDX-MS is not reliant on successful crystallization trials, we saw this as an opportunity to describe the solution structural integrity of apo-USP30, which would allow us to elucidate its mode of binding to **39**.

The conformational landscape of apo-USP30 generally follows the arrangement of its USP domains; some of the most solvent exposed regions of the protein are found at the linker regions connecting individual domains, most noticeably between USP domains 1 and 2, 4 and 5, and 5 and 6 (**Figures S3a** and **S3b**). Conversely, USP domains 1 and 5 and the N-terminal end of USP domain 6 are largely protected from the solvent and inaccessible (**Figure S3b**). This is in good agreement with HDX-MS data recently acquired on the full length apo-USP30 protein, where the USP domains were shown to be in a conformation that was generally hidden from the solvent and connected by several exposed linkers.^10^ However, due to the instability of the full-length species, HDX-MS was performed on a much smaller scale compared to our own study, with only a sole 3 sec labeling time point measured. Furthermore, an appreciation of USP30 dynamics could not be extracted from this single time point. Looking at apo-USP30 dynamics across the several labeling time points described herein, an increase in the rate of deuteration over the time course of the experiment was observed across the majority of the protein (**Figure S3b**). This dynamic HDX-MS behavior is indicative of the presence of secondary structural elements, thereby confirming the highly structured nature of the protein. The regions with the greatest dynamic HDX-MS behavior were found within USP domains 2, 3, 4, and 6 (**Figure S3b**). Conversely, no dynamic HDX-MS events were observed in several regions of apo-USP30 and the maximum level of deuteration was reached immediately, indicating structural disorder. These unstructured regions map to the N- and C-terminal extremities of the protein and within USP domains 1 (residues 71-78), 2 (residues 130-136), and 5 (residues 439-453). There is a high level of overlap when mapping the structural data inferred from the HDX-MS to the crystal structure of 5OHK for both apo- and holo-USP30 (**Figure S4**).

Several regions of USP30 undergo structural transitions in the presence of **39**, which are potentially significant in terms of inhibitory mechanistics. First, multiple segments of USP30 become completely blocked and inaccessible to the solvent following inhibitor binding. These include a region directly adjacent to the catalytic Cys77 at F78-L83 and an area of USP domain 2 covering E158-F166, as highlighted in purple in **Figure S4**. This suggests that **39** induces a conformation of USP30 that not only blocks off the catalytic region and its surroundings from the solvent, but importantly, also prevents access and binding of Ub itself. A second structural phenomenon is also evident, specifically the conversion of intrinsically disordered loops in the absence of **39**, to rigid, structural elements in the presence of the compound (**Figure S4**). These disorder-to-order transitions likely embody functional significance^31^ and in USP30, are found at the catalytic Cys77, represented by peptide V71-F78, a section of USP domain 2 at R148-F154, and a long chain of residues spanning Q326-L349.

Tracking these structural rearrangements across individual labeling time points allows us to propose a general timeline of inhibition (**Figure S5**). Taking Q326-L349 as an example, peptides mapping to this region of USP30 undergo significant structural transitions at the earliest time points monitored (30 and 60 sec), which are completed in the later stages of our experimental time course. Conversely, peptides proximal to the catalytic Cys77 become blocked and solvent inaccessible primarily in the latter half of our experiment (600 and 3600 sec). The fact that the residue (and adjacent regions) most crucial to USP30 catalysis, Cys77, is most significantly perturbed in the latter stages of our experiment could go some way to explain the slow and tight binding behavior observed for **39** in our enzyme kinetics analyses (**Figure 2**).

### Molecular docking proposes key residues important for 39 binding to USP30

To further refine our HDX-MS findings, we explored the binding mode of **39** to USP30 computationally. We performed molecular docking simulations using the simple docking mode in AMDock software, with the human USP30 catalytic domain from the crystal structure of USP30 in complex with ubiquitin propargylamide (Ub-PA; PDB code = 5OHK) acting as the target receptor for the compound.^10, 19, 32^ The *in silico* binding pose of **39** with the highest-ranking docking score has estimated affinity and Ki values of −7.9 kcal/mol and 1.62 μM respectively, compared with a reported experimental IC_50_ value of approximately 20 nM.^19^ Compound **39** is predicted to bind to the thumb-palm cleft that guides the Ub C-terminus into the active site, residing approximately 7.4 Å away at its closest point from the thiol side chain of catalytic Cys77 (**Figure 4a**). The benzyl moiety of **39** is flanked by Pro336, Met448, and the side chains of Leu328 and His449 (**Figure 4b**). The fluorophenyl moiety is flanked by the side chains of Leu328, Arg327, Lys338, and Tyr495, with a π-stacking interaction with His444. The N-*tert*-butyl sulfonamide moiety is anchored by hydrogen-bonding interactions with the side chains of Gln326 and His163. Compared with the structure of USP30 in complex with Ub-PA (PDB code = 5OHK), the modelled position of **39** would sterically clash with the C-terminal tail of the Ub substrate with the fluorophenyl moiety sitting in an equivalent position to the side chain of Ub Leu73, thereby preventing Ub binding and isopeptide bond cleavage (**Figure 4c**).

**Figure 4.**
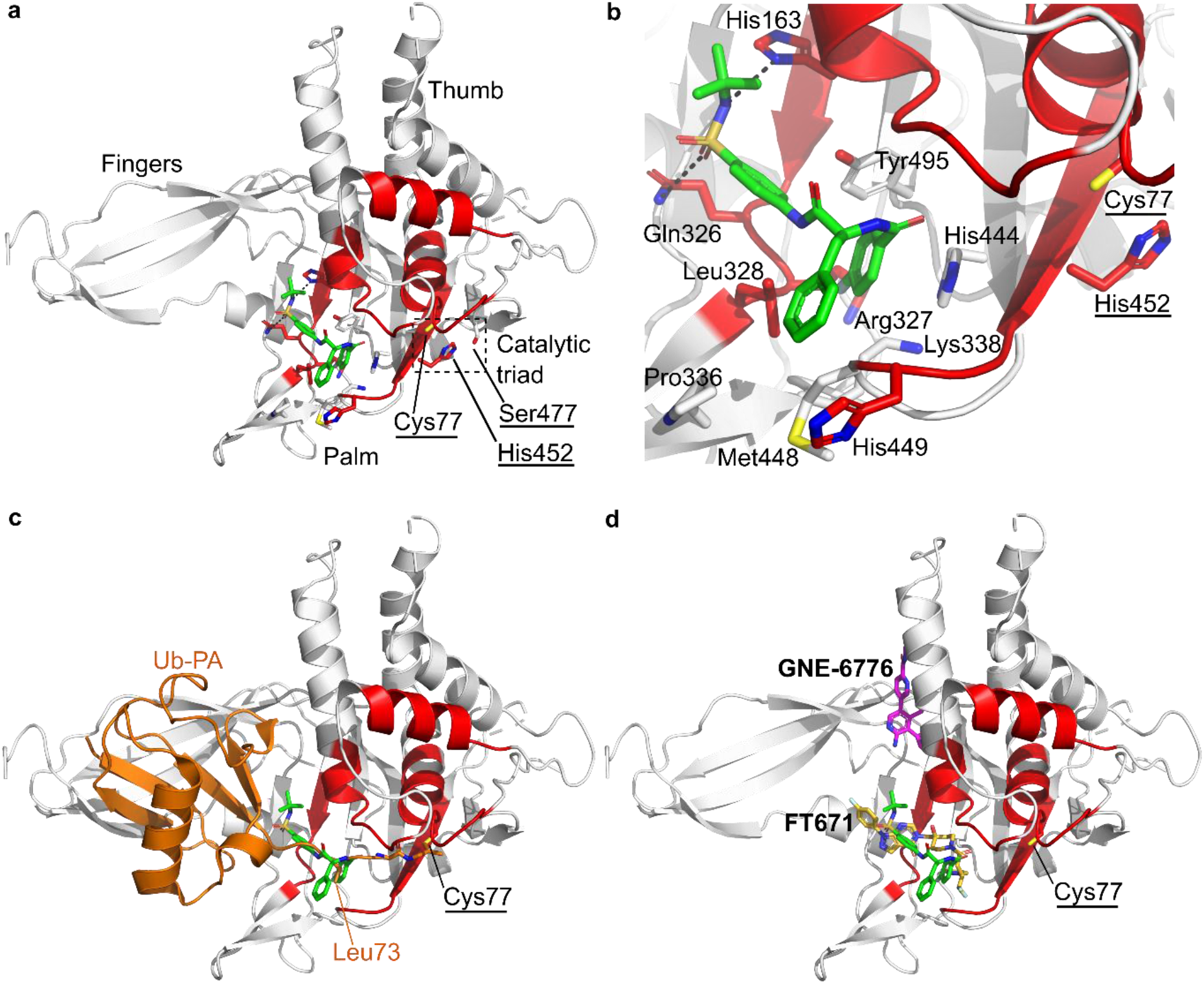
Modelled structure of human USP30 in complex with 39. (a) Structure of human USP30 catalytic domain highlighting the modelled position of **39** shown as a stick representation and colored green. The thumb, palm and fingers subdomains of the catalytic domain and catalytic triad (Cys77, Ser477 and His42; underlined) are highlighted. Regions identified in the HDX-MS analysis of USP30 in the presence of **39** are colored red. (b) Close-up view of the putative **39** binding site highlighting key residues and hydrogen-bonding interactions represented as dotted lines. (c) Superposition of Ub-PA (orange; PDB code= 5OHK) on the docked structure. **39** sterically clashes with the C-terminal tail of the Ub substrate thereby preventing Ub binding and isopeptide bond cleavage. (d) Superposition of the USP7 inhibitors, FT671 (yellow carbon atoms) and GNE-6776 (magenta carbon atoms) in complex with USP7, onto the docked structure. **39** putatively binds to an equivalent site in the thumb-palm cleft compared with FT671. Figure prepared using PyMOL (The PyMOL Molecular Graphics System, Version 2.4.1, Schrödinger, LLC.).

Crystal structures of USP7 in complex with the small-molecule inhibitors, FT761 (PDB code= 5NGE)^30^ and GNE-6776 (PDB code= 5UQX)^33^, reveal two distinct inhibitor binding modes that attenuate Ub binding and inhibit the DUB activity. FT671 binds to the thumb-palm cleft and resides approximately 5 Å away from the catalytic cysteine whereas GNE-6776 interacts with acidic residues in the USP7 catalytic domain that mediate hydrogen-bonding interactions with the Ub Lys48 side chain and binds 12 Å from the catalytic cysteine. A comparison of the docked structure of **39** with the USP7 inhibitor complexes suggests that **39** is most likely to bind to an equivalent site to FT671 (**Figure 4d**). In addition, the docked pose of **39** correlates well with the HDX-MS data, with the predicted binding site of **39** being flanked by residues residing in peptides E158-F166, I323-L328, and H449-Y456, which become solvent protected in HDX-MS (**Figure 4a** and **b**). The HDX-MS analysis also implicates peptides N72-L83 (which contains the catalytic Cys77) and D139-M149 in structural rearrangements upon compound binding. These regions reside further from the predicted binding site of **39**. However, compound binding may potentially cause conformational rearrangements of the catalytic domain remote from the binding site resulting in these regions becoming protected upon compound binding. Similar conformational rearrangements are seen in human USP7, in which the inhibitors bind to the catalytically incompetent apoform state with switching loop “in” as compared with the catalytically competent Ub-bound state with switching loop “out”, and it is possible that USP30 may exhibit similar dynamic conformational flexibility.

## CONCLUSIONS

Mitochondrial pathway disruption has been linked to a spectrum of pathophysiological conditions, from neurodegeneration and acute, chronic kidney and cardiovascular diseases, through to hepatocellular carcinoma and peroxisome biogenesis disorders.^6, 34–36^ USP30 represents an actionable drug target of these conditions through its participation in PINK1/Parkin-mediated mitophagy, BAX/BAK-dependent apoptosis, oncogenesis, and pexophagy.^18, 37–39^ USP30 regulates mitophagy by antagonizing Parkin-mediated ubiquitination and its inhibition has been shown to have significant therapeutic potential against PD and similar neurodegenerative disorders. Drug discovery efforts targeting USP30 have yielded the highly potent and selective small-molecule benzosulfonamide inhibitor, compound **39**.^17, 19, 40^ Combining state-of-the-art proteomics, HDX-MS and molecular docking, we have described the dynamic structural interplay between USP30 and **39**. The inhibitor binds USP30 in a slow and tight manner, and displays kinetic properties consistent with covalent attachment to USP30, despite its non-covalent design. Collectively, our integrative structural biology lens successfully identified regions within USP30 that undergo dramatic structural and conformational rearrangements in the presence of **39**, which prevent Ub binding and decrease DUB activity. X-ray data for USP30 in complex with **39** will undoubtedly complement these observations and will, combined with molecular dynamics studies, drive the development of next-generation inhibitors.

## EXPERIMENTAL SECTION

### PURITY

All compounds are >95% pure by HPLC analysis (**Figure S6**).

#### 1. ABPP ASSAY

##### HA-Ub-PA synthesis

HA-Ub-PA synthesis was carried out as previously described.^27, 41^ Briefly, a pTYB construct was used to express Ub (Gly76del) in *E.coli*. The Ub was tagged with a HA tag on the N-terminus, and an intein-chitin binding domain on the C terminus. *E.coli* lysis was performed using sonication in 50 mM Hepes, 150 mM NaCl, 0.5 mM Dithiothreitol (DTT). The protein was then bound to chitin bead slurry and incubated with 100 mM MesNa overnight (37°C with agitation) to form HA-Ub-MesNa. The HA-Ub-MesNa was then incubated for 20 min with 250 mM propargylamine (PA; room temperature with agitation), and desalted to remove excess propargylamine, resulting in the reactive activity-based probe HA-Ub-PA.

##### Cell culture and lysis

SH-SY5Y cells were cultured at 37°C, 5% CO_2_, in Eagle’s Minimum Essential Medium and Ham’s F12 Nutrient Mix (1:1), supplemented with 15% Fetal Bovine Serum, 1% non-essential amino acids and 2 mM Glutamax. Cells were collected by washing with phosphate-buffered saline (PBS), followed by scraping in PBS and centrifugation at 200 x g. Cells were lysed in 50 mM Tris Base, 5 mM MgCl_2_.6 H_2_O, 0.5 mM EDTA, 250 mM Sucrose, 1 mM DTT by vortexing with acid washed beads (1:1 v/v) 10 times (30 seconds vortexing, 1 min break on ice). Lysates were clarified at 600 x g for 10 min at 4°C. Lysate protein concentration was then determined by BCA.

##### Inhibitor selectivity with HA-Ub-PA immunoprecipitation

HA-Ub-PA protein complexes were immunoprecipitated and analysed using LC-MS/MS as previously described.^42^ Inhibitor **39** or DMSO was incubated with 500 μg of SH-SH5Y protein lysate for 1 h at 37°C. HA-Ub-PA was then incubated with the **39**-treated lysates for 45 min at 37°C at a protein ratio of 1:200 (w/w). The reaction was quenched with the addition of 0.4% SDS and 0.5% NP-40 (IGEPAL CA-630) and diluted to 0.5 mg/mL with 50 mM Tris, 0.5% NP-40, 150 mM NaCl and 20 mM MgCl_2_.6 H_2_O, pH 7.4. HA-Ub-PA protein complexes were then immunoprecipitated using 150 μL of pre-washed Anti-HA agarose slurry overnight at 4°C with end-over-end rotation. The agarose slurry was then washed four times and the HA-Ub-PA protein complexes were eluted using 110 μL of 2 x Laemmli buffer. To check for efficient immunoprecipitation, 10 μL of the eluates were run on a western blot.

The remaining 100 μL of the eluates were reduced with 20 mM DTT for 10 min at 95°C and alkylated with 40 mM of iodoacetamide for 30 min at room temperature in the dark. Proteins were then acidified to 1.2% phosphoric acid, diluted 6-fold with 90% methanol/100 mM TEAB, and captured/washed on an S-trap column according to the standard protocol.^43^ Proteins were digested on the S-trap column with 2 μg of trypsin overnight at 37°C. Eluted peptides were then dried and resuspended in 2% acetonitrile (ACN), 0.1% formic acid.

##### LC-MS/MS

Peptides were analysed using a Dionex Ultimate 3000 nano-ultra high pressure reversed-phase chromatography system coupled on-line to an Orbitrap Fusion Lumos mass spectrometer (Thermo Scientific). Samples were separated on an EASY-Spray PepMap RSLC C18 column (500 mm x 75 μm, 2 μm particle size; Thermo Scientific) over a 60 min gradient of 2-35% ACN in 5% DMSO, 0.1% formic acid and at 250 nL/min. The column temperature was maintained at 50°C with the aid of a column oven. The mass spectrometer was operated in positive polarity mode with a capillary temperature of 275°C. Data-independent acquisition (DIA) mode was utilized for automated switching between MS and MS/MS acquisition as described previously.^23^ Fractions were loaded with adjusted sample volumes to analyze ~200 ng on column.

##### DIA-MS data processing and analysis

Data was analysed using DIA-NN (version 1.8) with all settings as default. A *Homo sapiens* Uniprot database (retrieved on 16/04/2021) was used for the analysis. Immunoprecipitations were carried out in duplicate and any DUBs that were not present in both control samples and enriched > 5-fold when compared to a no probe control, were discarded from the analysis. Identifications that were assigned to multiple DUBs were not included in the analysis. MINDY3 was also removed from the dataset as it was the lowest intensity, so may have been at the bottom of the instruments dynamic range and did not produce stable values across the dataset. All MS raw files were deposited in PRIDE under the code PXD036574.

#### 2. ENZYME KINETICS

##### In Vitro USP30 Activity Assay

Fluorescence intensity measurements were used to monitor the cleavage of a Ub-rhodamine (Ub-Rho110) substrate. All activity assays were performed in black 384-well plates in 20 mM Tris-HCl, pH 8.0, 150 mM Potassium Glutamate, 0.1 mM TCEP and 0.03% Bovine Gamma Globulin with a final assay volume of 20 μL. Compound IC_50_ values for DUB inhibition were determined as previously described.^17^ Briefly, an 11-point dilution series of compounds were dispensed into black 384-well plates using an Echo-550 Acoustic Liquid Handler (Beckman Coulter). USP30, 0.2 nM (residues 64-502Δ179-216 & 288-305, Viva Biotech (Shanghai) Ltd.) was added and the plates preincubated for 30 min, 25 nM Ub-Rho110 (Ubiquigent) was added to initiate the reaction and the fluorescence intensity was recorded for 30 min on a Pherastar FSX (λ_Ex_ = 485 nm, λ_Em_ = 520 nm) (BMG Labtech). Initial rates were plotted against compound concentration to determine IC_50_. Data was processed using analysis tools from Dotmatics (https://www.dotmatics.com/).

##### Kinetic assays - determination of kinetic parameters for slow-tight binding inhibitors

Kinetic assays were performed in 384-well Sensoplate™ in 20 mM Tris-HCl, pH 8.0, 300 mM Potassium Glutamate, 0.1 mM TCEP and 0.2% BGG with a final assay volume of 50 μL. An 11-point dilution series of compound was dispensed into assay plates and 25 μL 2X Ub-Rho110 was added. The dispense function of the FLIPR^®^ Tetra (Molecular Devices) was used to add 25 μL 2X USP30 to give final assay concentrations of 5 and 180 nM for USP30 and Ub-Rho110 respectively. The fluorescence signal of the enzyme activity was monitored every 3 sec for 1800 sec (λ_Ex_ = 470-495 nm, λ_Em_ = 515-575 nm, camera gain 70, exposure time 0.6 sec, excitation intensity 80%). Analysis was performed in GraphPad Prism version 9.4.1 for Windows (Graphpad Software, La Jolla, California, USA; www.graphpad.com). The time course data was normalized relative to enzyme in the absence of compound and used to generate inhibition curves at each time point.

**Scheme A.**
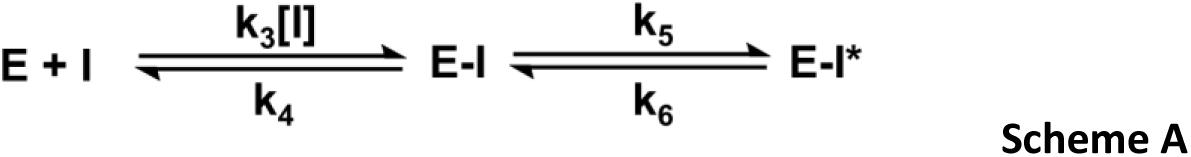

As IC_50_ values are time-dependent for compound **39** with no covalent labeling of USP30, shown by MS (**Figure S1**), data were modelled to a slow-tight binding scheme (**Scheme A**). Fitting of progress curves allows for calculation of relevant kinetic parameters (**Supporting Information**).

##### Bio-layer interferometry

Bio-layer interferometry was performed on an Octet RED384^®^ system (Sartorius) at 25°C in a buffer containing 20 mM Tris-HCl (pH 8), 100 mM NaCl, 2 mM TCEP, 0.05% Tween and 1% DMSO. Biotinylated USP30 (residues 64-502Δ179-216 & 288-305, Viva Biotech (Shanghai) Ltd.) was immobilized onto Super Streptavidin (SSA) biosensors. After 60 sec baseline detection, the association of defined concentrations of **39** (0-5 μM) was recorded over 300 sec followed by dissociation in buffer over 600 sec. Traces were normalised by double subtraction of baseline (no compound) and reference sensors (no USP30, association and dissociation of compound) to correct for non-specific binding to the sensors. Traces were analysed using the Octet Software (Version 11.2, Sartorius).

#### 3. HDX-MS

##### HDX sample preparation

USP30 was incubated in either the presence (holo-USP30) or absence (apo-USP30) of a two-fold molar excess of **39**, ensuring that all complexes were fully formed and maintained over the course of the labeling reaction. Before the HDX-MS experiments, labelling (L), equilibration (E) and quench (Q) buffers were freshly prepared with D_2_O or H_2_O, respectively (Buffers E and L: 50 mM HEPES, 400 mM NaCl, 2.0 mM TCEP, 10% glycerol (v/v) at pH 7.2; Buffer Q: 50 mM potassium phosphate buffer, 2.0 M guanidine hydrochloride at pH 2.30). The USP30 protein sample was supplied at 66 μM and was diluted in Buffer E to a final concentration of 11 μM, which equates to 16 pmol injected onto the pepsin column. Buffers E and L were equilibrated at 20°C, while the protein samples and Buffer Q were equilibrated at 0°C.

##### HDX cyclic ion mobility mass spectrometry

HDX-MS experiments were carried out on a fully automated HDX-2 system (supplied by Waters, Milford USA) previously described by Brown and Wilson.^44^ The exchange reaction was initiated by diluting 3.5 μL protein sample with a concentration of 11 μM into 56.5 μL Buffer E for reference, or Buffer L for D_2_O labeling, and incubated for several time points (0, 30, 60, 600 and 3600 sec). A D_2_O/H_2_O ratio in excess of 90% guaranteed that the kinetics favored unidirectional exchange. Subsequently, the exchange reaction was stopped by mixing 50 μL of sample with 50 μL pre-cooled Buffer Q. Next, 50 μL of quenched sample was subjected to a temperature-controlled chromatography system (HDX M-Class UPLC, Waters). The protein was digested online by a pepsin column (Enzymate BEH pepsin column; 2.1 x 30 mm; Waters). Eluting peptides were trapped and washed on a C18 pre– column (C18 1.7 μM VanGuard 2.1 x 5 mm pre-column; Waters) at 100 μL/min for 3 min and separated on a reversed phase column (C18 1.7 μM Acquity UPLC 1 x 100 mm reverse phased column; Waters) with a linear gradient ranging from 5% ACN to 40% ACN plus 0.2% FA at 40 μL/min in 8 min, followed by a rapid rise to 99% ACN holding for 0.3 min. ACN concentration was rapidly reduced to 5% and held there for 0.2 min, followed by a linear gradient back to 99% over 0.7 min, and holding that concentration for 0.1 min. Next, C18 columns were equilibrated with 95% H_2_O plus 0.2% FA for 4 min. The reversed-phase chromatographic system was kept at approximately 0°C to reduce back-exchange. Peptides eluting from reversed phase column were measured with a SELECT SERIES Cyclic IMS mass spectrometer (Waters, Wilmslow UK) in HDMS^E^ mode (*m/z* 50-2000). This mode utilises ion mobility (IM) separation for orthogonal separation of the peptides (LC, IM, *m/z*). The mass spectrometer was fitted with an electrospray source equipped with additional independent LockSpray probe (Leu-enkephalin LockMass solution was used, *m/z* 556.2771).

##### HDX-MS data processing and analysis

All MS-analyses were performed in triplicate for each time point and condition. Protein Lynx Global Server (PLGS) 3.0 (Waters Corporation) was used for peptide identifications. A peptic peptide sequence coverage map was generated in DynamX 3.0 HDX software (Waters Corporation). Peptide-level deuterium uptake data was also visualized in DynamX (**Figure S7**) and reported as relative deuterium exchange levels expressed in either mass unit or fractional exchange. The latter was calculated by dividing the experimentally measured uptake by the theoretically maximum number of exchangeable backbone amide hydrogens that could be replaced within each peptide. This number corresponds to the number of amino acid residues present in the peptide minus the number of proline residues and minus one for the N-terminus that back exchanges too rapidly to be measured by MS.^31^ A single charge state was considered per peptide. Data were also verified and visualized in MEMHDX.^45^

#### 4. MOLECULAR DOCKING

The crystal structure of human USP30 catalytic domain (residues K64-V502) in covalent complex with Ub-PA at 2.34 Å resolution represents the highest resolution human USP30 structure available in the Protein Data Bank (PDB code= 5OHK)^10^, and was used as the target receptor for docking the selective USP30 benzosulfonamide inhibitor, compound **39** ^19^, using AutoDock Vina implemented in the program AMDock 1.5.2.^32^ Coordinates for **39** were generated using ChemDraw Prime 16.0.1.4, and PRODRG implemented in the CCP4 software suite.^46^ The compound was docked using the simple docking mode and automatic defined search space in AMDock 1.5.2.

## ANCILLARY INFORMATION

### Supporting Information

This can be accessed here xxx

### PDB ID Codes

*5OHK* represents the crystal structure of human USP30 catalytic domain (residues K64-V502) in covalent complex with Ub propargylamide (Ub-PA) at 2.34 Å resolution. This is the highest resolution structure of human USP30 currently available in the PDB.

### Homology Models

Not applicable

### Corresponding Author Information

Dr Darragh P O’Brien can be contacted at

Professor Benedikt M Kessler can be contacted at

### Present/Current Author Addresses

*Target Discovery Institute, Centre for Medicines Discovery, Nuffield Department of Medicine, University of Oxford, UK* - Darragh P O’Brien, Hannah BL Jones, Benedikt M Kessler

*ARUK-Oxford Drug Discovery Institute, Centre for Medicines Discovery, Nuffield Department of Medicine, University of Oxford, UK* - Franziska Guenther, Katherine S England, Emma J Murphy, Paul Brennan, John B Davis

*Waters Corporation, Wilmslow, Cheshire, UK* - Malcolm Anderson

*Chinese Academy of Medical Sciences – Oxford Institute, Nuffield Department of Medicine, University of Oxford, UK* - Adán Pinto-Fernández, Benedikt M Kessler

*Cancer Research Horizons, Francis Crick Institute, London, UK* - Andrew P Turnbull

### Author Contributions

JD and BMK designed the study. DPOB, HBLJ, FG, EM, MA, APT, and BMK wrote the first manuscript draft. DPOB, HBLJ, FG, EM, MA, and APF designed and performed the experiments. APT designed and performed the molecular docking. All authors discussed the results and commented on the manuscript.

## Acknowledgments

We wish to express thanks to Drs Roman Fischer and Iolanda Vendrell from the Oxford Discovery Proteomics Facility for their help with ABPP-MS data acquisition. We would like to thank Daryl S Walter from Evotec and Jeff Schkeryantz from Bristol Myers Squibb for their highly useful project discussions. Our gratitude goes to Alzheimer’s Research UK for their support and funding for the ARUK-Oxford Drug Discovery Institute (grant no. ARUK-2021DDI-OX). We would also like to thank our other funders, including the Chinese Academy of Medical Sciences (CAMS) Innovation Fund for Medical Science (CIFMS), China (grant number: 2018-I2M-2-002) (BMK, APF), Bayer (BMK), the late Mr & Mrs James Hardwick for funding the ODDI Medicinal Chemistry Team and also the G & K Boyes Charitable Trust.

## Abbreviations Used

Ub: Ubiquitin
UPS: Ubiquitin Proteasome System
PD: Parkinson’s Disease
MOM: Mitochondrial Outer Membrane
SAR: Structure Activity Relationship
USP30: Ubiquitin Specific Protease 30
DUB: Deubiquitinase
ABPP-MS: Activity-Based Protein Profiling Mass Spectrometry
LFQ: Label-Free Quantitation
DIA: Data Independent Acquisition
HDX-MS: Hydrogen Deuterium eXchange Mass Spectrometry
PDB: Protein Data Bank

## REFERENCES

1. Popovic, D.; Vucic, D.; Dikic, I., Ubiquitination in disease pathogenesis and treatment. Nat Med 2014, 20 (11), 1242–53.

2. Ciechanover, A., The ubiquitin proteolytic system and pathogenesis of human diseases: a novel platform for mechanism-based drug targeting. Biochem Soc Trans 2003, 31 (2), 474–81.

3. Pickart, C. M.; Eddins, M. J., Ubiquitin: structures, functions, mechanisms. Biochim Biophys Acta 2004, 1695 (1–3), 55–72.

4. Mulder, M. P.; Witting, K.; Berlin, I.; Pruneda, J. N.; Wu, K. P.; Chang, J. G.; Merkx, R.; Bialas, J.; Groettrup, M.; Vertegaal, A. C.; Schulman, B. A.; Komander, D.; Neefjes, J.; El Oualid, F.; Ovaa, H., A cascading activity-based probe sequentially targets E1-E2-E3 ubiquitin enzymes. Nat Chem Biol 2016, 12 (7), 523–30.

5. Pollock, L.; Jardine, J.; Urbe, S.; Clague, M. J., The PINK1 repertoire: Not just a one trick pony. Bioessays 2021, 43 (11), e2100168.

6. Bingol, B.; Sheng, M., Mechanisms of mitophagy: PINK1, Parkin, USP30 and beyond. Free Radic Biol Med 2016, 100, 210–222.

7. Komander, D.; Clague, M. J.; Urbe, S., Breaking the chains: structure and function of the deubiquitinases. Nat Rev Mol Cell Biol 2009, 10 (8), 550–63.

8. Nguyen, T. N.; Padman, B. S.; Lazarou, M., Deciphering the Molecular Signals of PINK1/Parkin Mitophagy. Trends Cell Biol 2016, 26 (10), 733–744.

9. Marcassa, E.; Kallinos, A.; Jardine, J.; Rusilowicz-Jones, E. V.; Clague, M. J.; Urbe, S., New aspects of USP30 biology in the regulation of pexophagy. Autophagy 2019, 15 (9), 1634–1637.

10. Gersch, M.; Gladkova, C.; Schubert, A. F.; Michel, M. A.; Maslen, S.; Komander, D., Mechanism and regulation of the Lys6-selective deubiquitinase USP30. Nat Struct Mol Biol 2017, 24 (11), 920–930.

11. Sato, Y.; Okatsu, K.; Saeki, Y.; Yamano, K.; Matsuda, N.; Kaiho, A.; Yamagata, A.; Goto-Ito, S.; Ishikawa, M.; Hashimoto, Y.; Tanaka, K.; Fukai, S., Structural basis for specific cleavage of Lys6-linked polyubiquitin chains by USP30. Nat Struct Mol Biol 2017, 24 (11), 911–919.

12. Pickrell, A. M.; Youle, R. J., The roles of PINK1, parkin, and mitochondrial fidelity in Parkinson’s disease. Neuron 2015, 85 (2), 257–73.

13. Agarwal, S.; Muqit, M. M. K., PTEN-induced kinase 1 (PINK1) and Parkin: Unlocking a mitochondrial quality control pathway linked to Parkinson’s disease. Curr Opin Neurobiol 2022, 72, 111–119.

14. Kitada, T.; Asakawa, S.; Hattori, N.; Matsumine, H.; Yamamura, Y.; Minoshima, S.; Yokochi, M.; Mizuno, Y.; Shimizu, N., Mutations in the parkin gene cause autosomal recessive juvenile parkinsonism. Nature 1998, 392 (6676), 605–8.

15. Valente, E. M.; Abou-Sleiman, P. M.; Caputo, V.; Muqit, M. M.; Harvey, K.; Gispert, S.; Ali, Z.; Del Turco, D.; Bentivoglio, A. R.; Healy, D. G.; Albanese, A.; Nussbaum, R.; Gonzalez-Maldonado, R.; Deller, T.; Salvi, S.; Cortelli, P.; Gilks, W. P.; Latchman, D. S.; Harvey, R. J.; Dallapiccola, B.; Auburger, G.; Wood, N. W., Hereditary early-onset Parkinson’s disease caused by mutations in PINK1. Science 2004, 304 (5674), 1158–60.

16. Yue, W.; Chen, Z.; Liu, H.; Yan, C.; Chen, M.; Feng, D.; Yan, C.; Wu, H.; Du, L.; Wang, Y.; Liu, J.; Huang, X.; Xia, L.; Liu, L.; Wang, X.; Jin, H.; Wang, J.; Song, Z.; Hao, X.; Chen, Q., A small natural molecule promotes mitochondrial fusion through inhibition of the deubiquitinase USP30. Cell Res 2014, 24 (4), 482–96.

17. Rusilowicz-Jones, E. V.; Jardine, J.; Kallinos, A.; Pinto-Fernandez, A.; Guenther, F.; Giurrandino, M.; Barone, F. G.; McCarron, K.; Burke, C. J.; Murad, A.; Martinez, A.; Marcassa, E.; Gersch, M.; Buckmelter, A. J.; Kayser-Bricker, K. J.; Lamoliatte, F.; Gajbhiye, A.; Davis, S.; Scott, H. C.; Murphy, E.; England, K.; Mortiboys, H.; Komander, D.; Trost, M.; Kessler, B. M.; Ioannidis, S.; Ahlijanian, M. K.; Urbe, S.; Clague, M. J., USP30 sets a trigger threshold for PINK1-PARKIN amplification of mitochondrial ubiquitylation. Life Sci Alliance 2020, 3 (8).

18. Wang, F.; Gao, Y.; Zhou, L.; Chen, J.; Xie, Z.; Ye, Z.; Wang, Y., USP30: Structure, Emerging Physiological Role, and Target Inhibition. Front Pharmacol 2022, 13, 851654.

19. Kluge, A. F.; Lagu, B. R.; Maiti, P.; Jaleel, M.; Webb, M.; Malhotra, J.; Mallat, A.; Srinivas, P. A.; Thompson, J. E., Novel highly selective inhibitors of ubiquitin specific protease 30 (USP30) accelerate mitophagy. Bioorg Med Chem Lett 2018, 28 (15), 2655–2659.

20. Rusilowicz-Jones, E. V.; Barone, F. G.; Lopes, F. M.; Stephen, E.; Mortiboys, H.; Urbe, S.; Clague, M. J., Benchmarking a highly selective USP30 inhibitor for enhancement of mitophagy and pexophagy. Life Sci Alliance 2022, 5 (2).

21. Engen, J. R.; Wales, T. E., Analytical Aspects of Hydrogen Exchange Mass Spectrometry. Annu Rev Anal Chem (Palo Alto Calif) 2015, 8, 127–48.

22. Marciano, D. P.; Dharmarajan, V.; Griffin, P. R., HDX-MS guided drug discovery: small molecules and biopharmaceuticals. Curr Opin Struct Biol 2014, 28, 105–11.

23. Demichev, V.; Messner, C. B.; Vernardis, S. I.; Lilley, K. S.; Ralser, M., DIA-NN: neural networks and interference correction enable deep proteome coverage in high throughput. Nat Methods 2020, 17 (1), 41–44.

24. Jones, H. B. L.; Heilig, R.; Davis, S.; Fischer, R.; Kessler, B. M.; Pinto-Fernandez, A., ABPP-HT*-Deep Meets Fast for Activity-Based Profiling of Deubiquitylating Enzymes Using Advanced DIA Mass Spectrometry Methods. Int J Mol Sci 2022, 23 (6).

25. Komander, D., Mechanism, specificity and structure of the deubiquitinases. Subcell Biochem 2010, 54, 69–87.

26. Clague, M. J.; Urbe, S.; Komander, D., Breaking the chains: deubiquitylating enzyme specificity begets function. Nat Rev Mol Cell Biol 2019, 20 (6), 338–352.

27. Jones, H. B. L.; Heilig, R.; Fischer, R.; Kessler, B. M.; Pinto-Fernandez, A., ABPP-HT - High-Throughput Activity-Based Profiling of Deubiquitylating Enzyme Inhibitors in a Cellular Context. Front Chem 2021, 9, 640105.

28. Krippendorff, B. F.; Neuhaus, R.; Lienau, P.; Reichel, A.; Huisinga, W., Mechanism-based inhibition: deriving K(I) and k(inact) directly from time-dependent IC(50) values. J Biomol Screen 2009, 14 (8), 913–23.

29. Cunningham, C. N.; Baughman, J. M.; Phu, L.; Tea, J. S.; Yu, C.; Coons, M.; Kirkpatrick, D. S.; Bingol, B.; Corn, J. E., USP30 and parkin homeostatically regulate atypical ubiquitin chains on mitochondria. Nat Cell Biol 2015, 17 (2), 160–9.

30. Turnbull, A. P.; Ioannidis, S.; Krajewski, W. W.; Pinto-Fernandez, A.; Heride, C.; Martin, A. C. L.; Tonkin, L. M.; Townsend, E. C.; Buker, S. M.; Lancia, D. R.; Caravella, J. A.; Toms, A. V.; Charlton, T. M.; Lahdenranta, J.; Wilker, E.; Follows, B. C.; Evans, N. J.; Stead, L.; Alli, C.; Zarayskiy, V. V.; Talbot, A. C.; Buckmelter, A. J.; Wang, M.; McKinnon, C. L.; Saab, F.; McGouran, J. F.; Century, H.; Gersch, M.; Pittman, M. S.; Marshall, C. G.; Raynham, T. M.; Simcox, M.; Stewart, L. M. D.; McLoughlin, S. B.; Escobedo, J. A.; Bair, K. W.; Dinsmore, C. J.; Hammonds, T. R.; Kim, S.; Urbe, S.; Clague, M. J.; Kessler, B. M.; Komander, D., Molecular basis of USP7 inhibition by selective small-molecule inhibitors. Nature 2017, 550 (7677), 481–486.

31. O’Brien, D. P.; Durand, D.; Voegele, A.; Hourdel, V.; Davi, M.; Chamot-Rooke, J.; Vachette, P.; Brier, S.; Ladant, D.; Chenal, A., Calmodulin fishing with a structurally disordered bait triggers CyaA catalysis. PLoS Biol 2017, 15 (12), e2004486.

32. Valdes-Tresanco, M. S.; Valdes-Tresanco, M. E.; Valiente, P. A.; Moreno, E., AMDock: a versatile graphical tool for assisting molecular docking with Autodock Vina and Autodock4. Biol Direct 2020, 15 (1), 12.

33. Kategaya, L.; Di Lello, P.; Rouge, L.; Pastor, R.; Clark, K. R.; Drummond, J.; Kleinheinz, T.; Lin, E.; Upton, J. P.; Prakash, S.; Heideker, J.; McCleland, M.; Ritorto, M. S.; Alessi, D. R.; Trost, M.; Bainbridge, T. W.; Kwok, M. C. M.; Ma, T. P.; Stiffler, Z.; Brasher, B.; Tang, Y.; Jaishankar, P.; Hearn, B. R.; Renslo, A. R.; Arkin, M. R.; Cohen, F.; Yu, K.; Peale, F.; Gnad, F.; Chang, M. T.; Klijn, C.; Blackwood, E.; Martin, S. E.; Forrest, W. F.; Ernst, J. A.; Ndubaku, C.; Wang, X.; Beresini, M. H.; Tsui, V.; Schwerdtfeger, C.; Blake, R. A.; Murray, J.; Maurer, T.; Wertz, I. E., USP7 small-molecule inhibitors interfere with ubiquitin binding. Nature 2017, 550 (7677), 534–538.

34. Honsho, M.; Okumoto, K.; Tamura, S.; Fujiki, Y., Peroxisome Biogenesis Disorders. Adv Exp Med Biol 2020, 1299, 45–54.

35. Schmidt, M. F.; Gan, Z. Y.; Komander, D.; Dewson, G., Ubiquitin signalling in neurodegeneration: mechanisms and therapeutic opportunities. Cell Death Differ 2021, 28 (2), 570–590.

36. Pan, W.; Wang, Y.; Bai, X.; Yin, Y.; Dai, L.; Zhou, H.; Wu, Q.; Wang, Y., Deubiquitinating enzyme USP30 negatively regulates mitophagy and accelerates myocardial cell senescence through antagonism of Parkin. Cell Death Discov 2021, 7 (1), 187.

37. Bingol, B.; Tea, J. S.; Phu, L.; Reichelt, M.; Bakalarski, C. E.; Song, Q.; Foreman, O.; Kirkpatrick, D. S.; Sheng, M., The mitochondrial deubiquitinase USP30 opposes parkin-mediated mitophagy. Nature 2014, 510 (7505), 370–5.

38. Liang, J. R.; Martinez, A.; Lane, J. D.; Mayor, U.; Clague, M. J.; Urbe, S., USP30 deubiquitylates mitochondrial Parkin substrates and restricts apoptotic cell death. EMBO Rep 2015, 16 (5), 618–27.

39. Riccio, V.; Demers, N.; Hua, R.; Vissa, M.; Cheng, D. T.; Strilchuk, A. W.; Wang, Y.; McQuibban, G. A.; Kim, P. K., Deubiquitinating enzyme USP30 maintains basal peroxisome abundance by regulating pexophagy. J Cell Biol 2019, 218 (3), 798–807.

40. Harrigan, J. A.; Jacq, X.; Martin, N. M.; Jackson, S. P., Deubiquitylating enzymes and drug discovery: emerging opportunities. Nat Rev Drug Discov 2018, 17 (1), 57–78.

41. Borodovsky, A.; Ovaa, H.; Kolli, N.; Gan-Erdene, T.; Wilkinson, K. D.; Ploegh, H. L.; Kessler, B. M., Chemistry-based functional proteomics reveals novel members of the deubiquitinating enzyme family. Chem Biol 2002, 9 (10), 1149–59.

42. Pinto-Fernandez, A.; Davis, S.; Schofield, A. B.; Scott, H. C.; Zhang, P.; Salah, E.; Mathea, S.; Charles, P. D.; Damianou, A.; Bond, G.; Fischer, R.; Kessler, B. M., Comprehensive Landscape of Active Deubiquitinating Enzymes Profiled by Advanced Chemoproteomics. Front Chem 2019, 7, 592.

43. HaileMariam, M.; Eguez, R. V.; Singh, H.; Bekele, S.; Ameni, G.; Pieper, R.; Yu, Y., S-Trap, an Ultrafast Sample-Preparation Approach for Shotgun Proteomics. J Proteome Res 2018, 17 (9), 2917–2924.

44. Brown, K. A.; Wilson, D. J., Bottom-up hydrogen deuterium exchange mass spectrometry: data analysis and interpretation. Analyst 2017, 142 (16), 2874–2886.

45. Hourdel, V.; Volant, S.; O’Brien, D. P.; Chenal, A.; Chamot-Rooke, J.; Dillies, M. A.; Brier, S., MEMHDX: an interactive tool to expedite the statistical validation and visualization of large HDX-MS datasets. Bioinformatics 2016, 32 (22), 3413–3419.

46. Winn, M. D.; Ballard, C. C.; Cowtan, K. D.; Dodson, E. J.; Emsley, P.; Evans, P. R.; Keegan, R. M.; Krissinel, E. B.; Leslie, A. G.; McCoy, A.; McNicholas, S. J.; Murshudov, G. N.; Pannu, N. S.; Potterton, E. A.; Powell, H. R.; Read, R. J.; Vagin, A.; Wilson, K. S., Overview of the CCP4 suite and current developments. Acta Crystallogr D Biol Crystallogr 2011, 67 (Pt 4), 235–42.

