## Supplementary Figures for "Structural Premise of Selective Deubiquitinase USP30 Inhibition by Small-Molecule Benzosulfonamides"

#### *Contents*

Figure S1: Confirmation of USP30 complex formation with Inhibitor 39.
Figure S2: Peptide map of USP30
Figure S3: Differential HDX-MS of USP30 and 39
Figure S4: Architecture of USP30 inferred by HDX-MS
Figure S5: Temporal exchange of USP30
Figure S6: HDX-MS uptake curves for individual USP30 peptides

**Supplementary Figure 1**

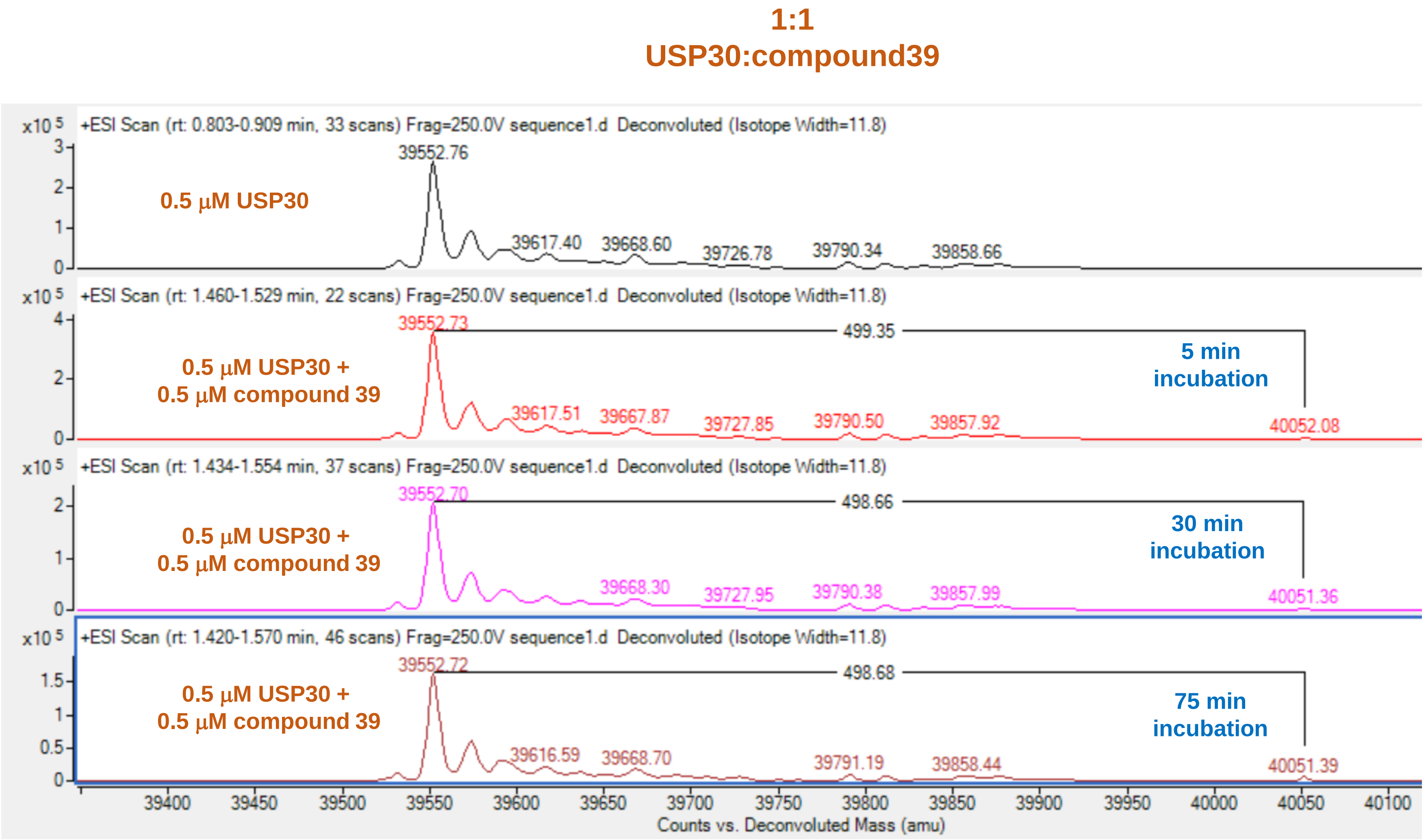


#### compound 39

**MW: 497.6**

#### USP30 (Viva)

- MW: **39552.24** Da
- insertion/deletion mutant: (64-178)-GSGS-(217-288)-G-(305-357) N346P, F348N, M350D, D352S, I353E, K355A, Y356S-SNA-(432-502)-6*His + biotin tag

### Supplementary Figure 2

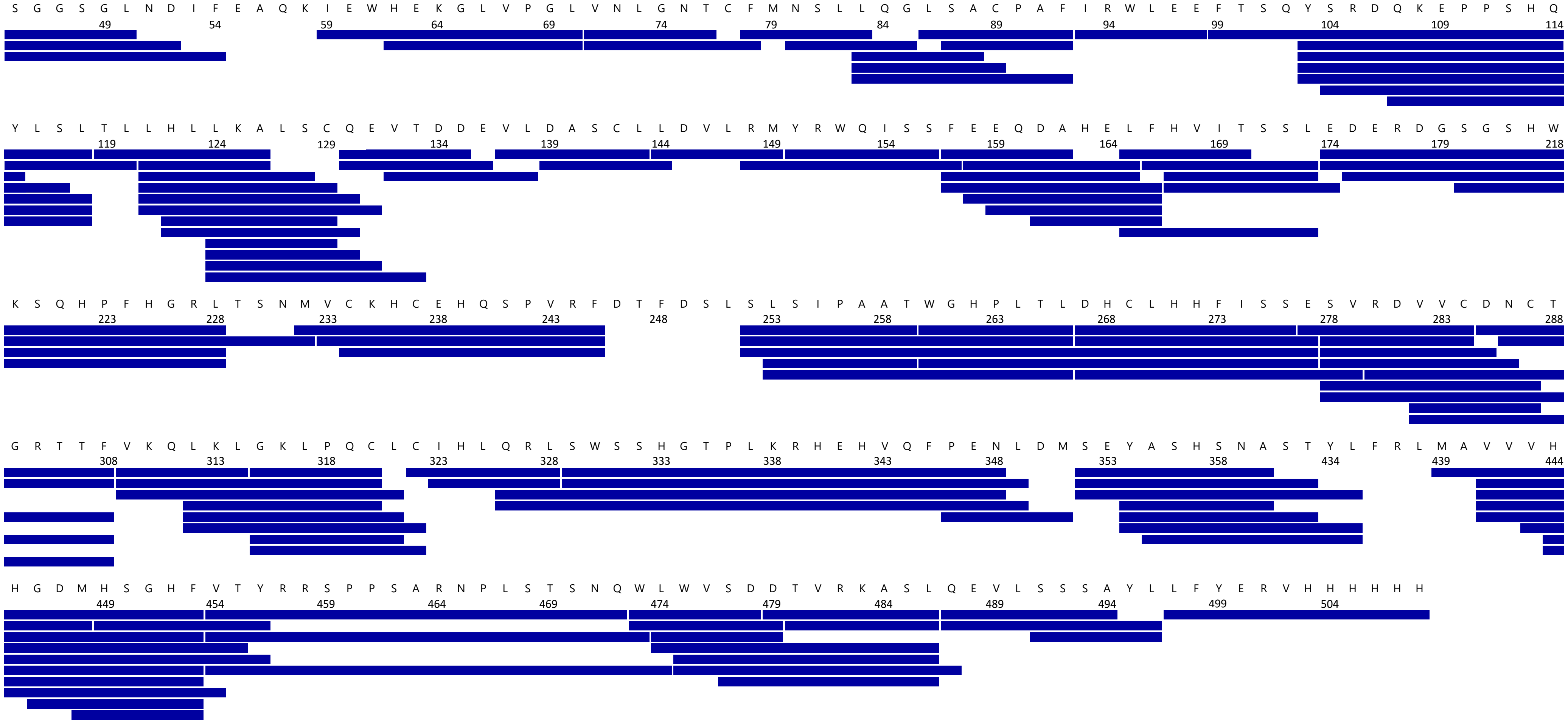

#### Supplementary Figure 3

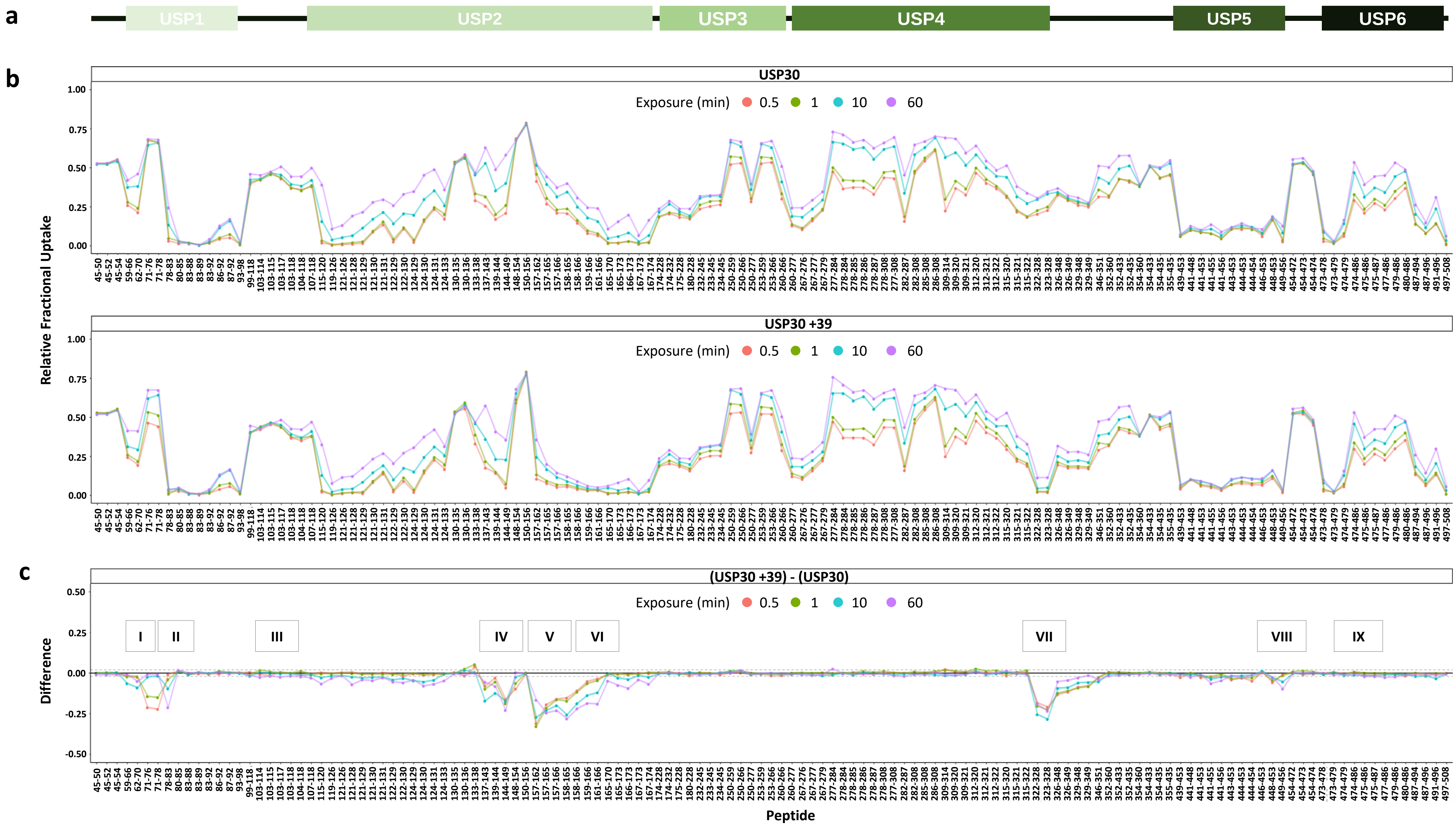

**Supplementary Figure 4**

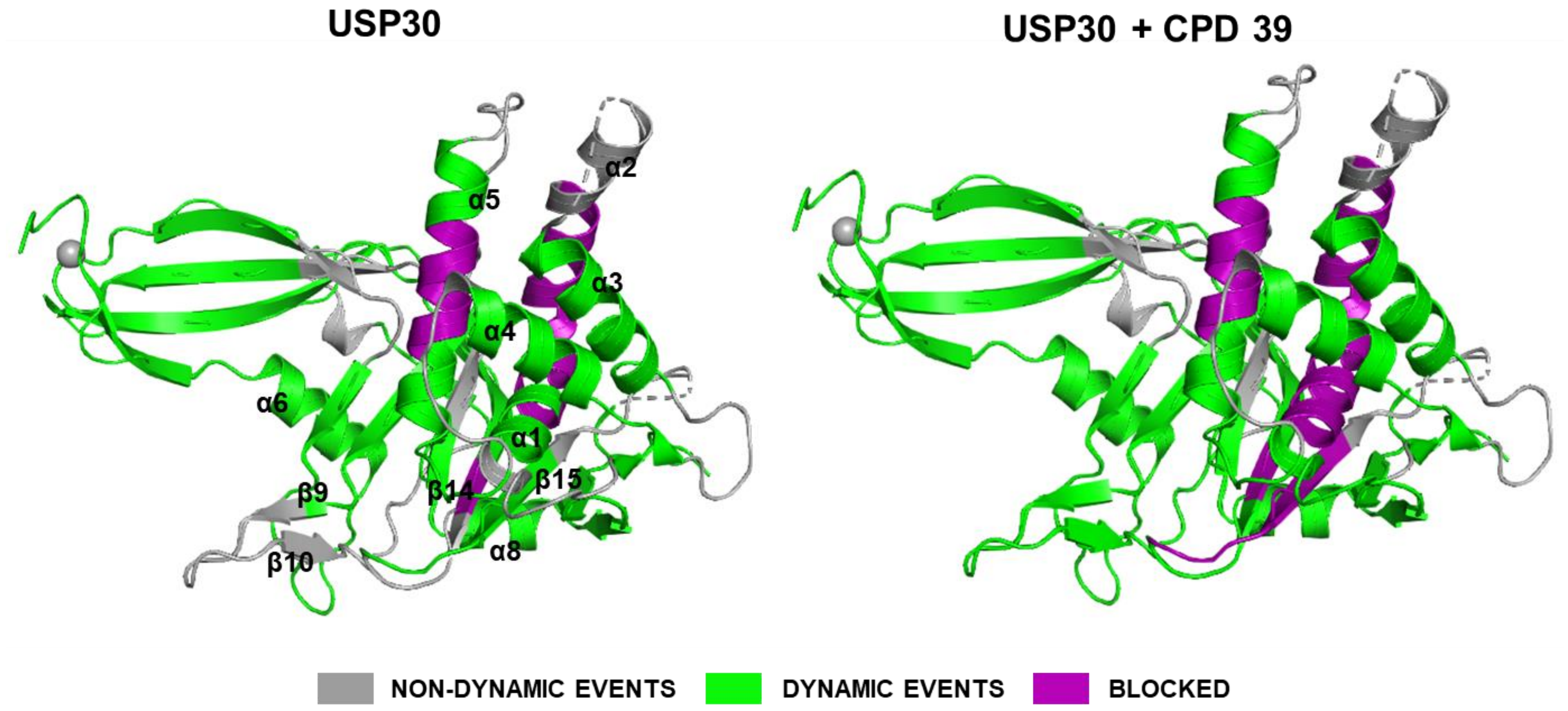

**Supplementary Figure 5**

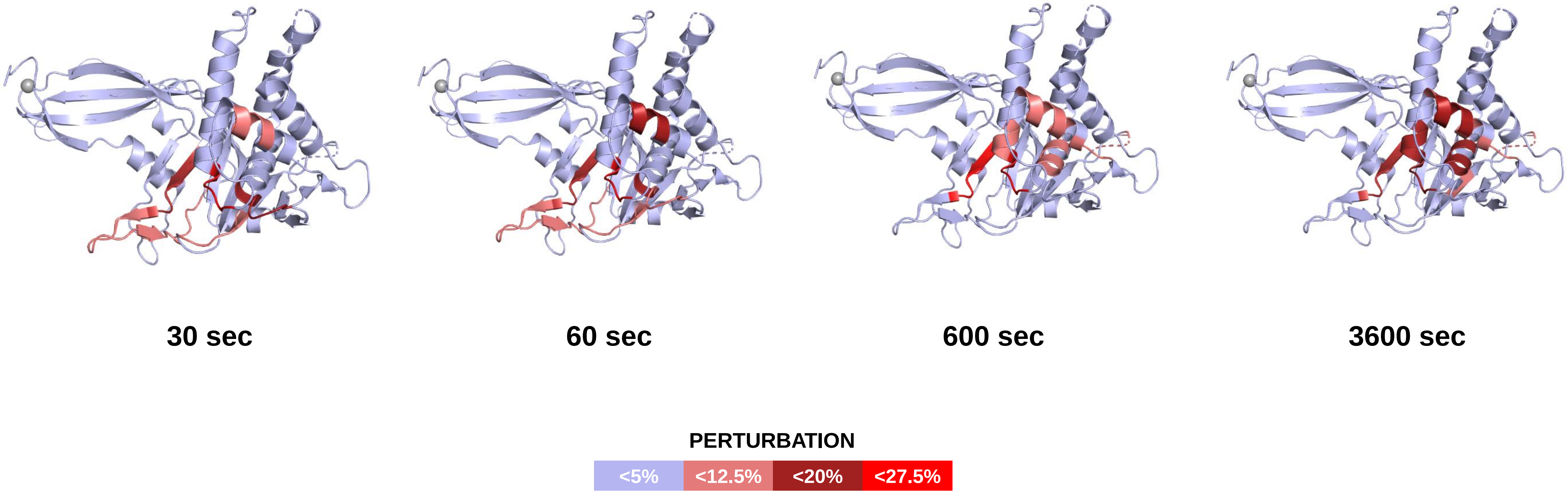

### Supplementary Figure 6

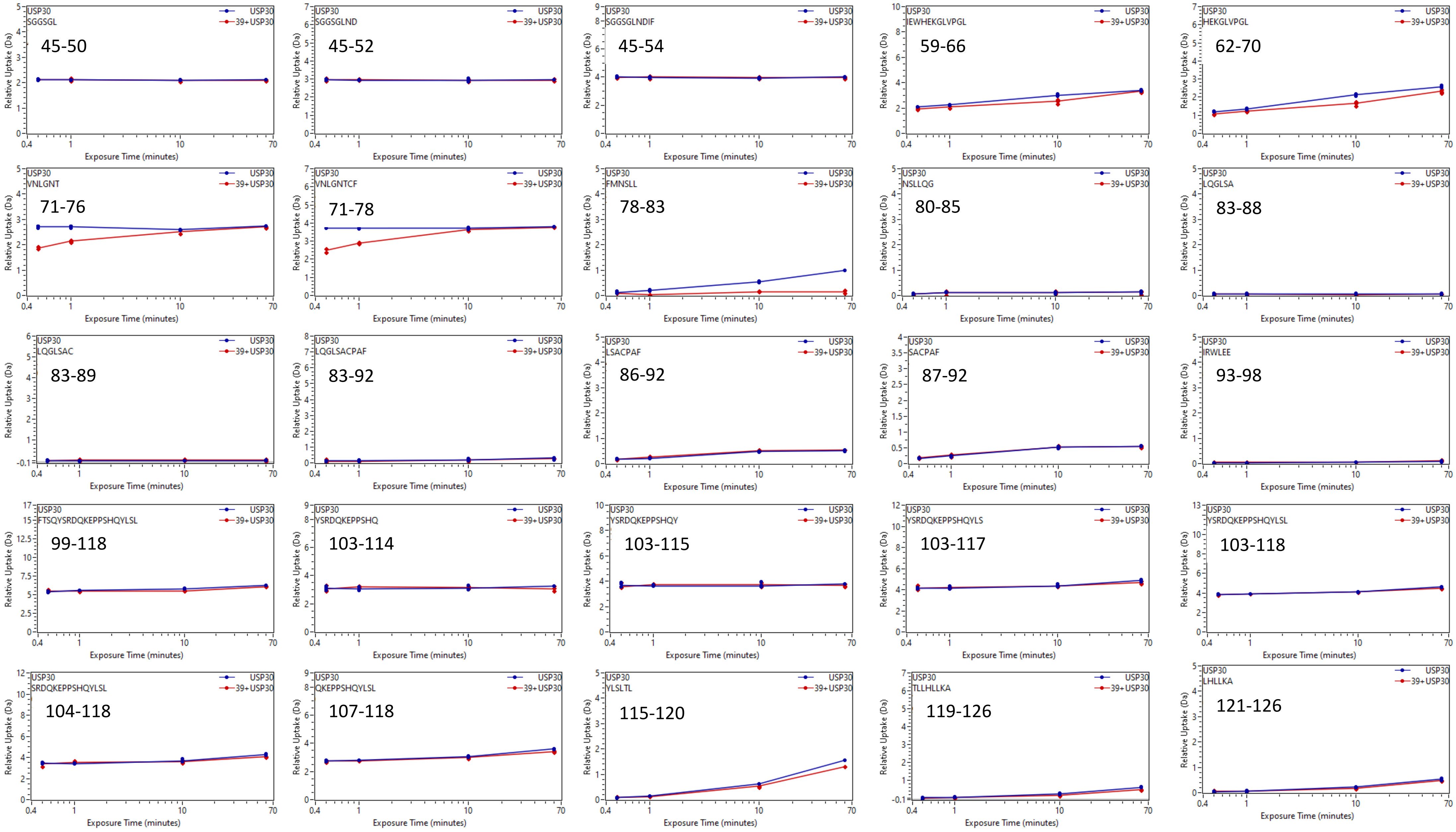

### Supplementary Figure 6 (continued)

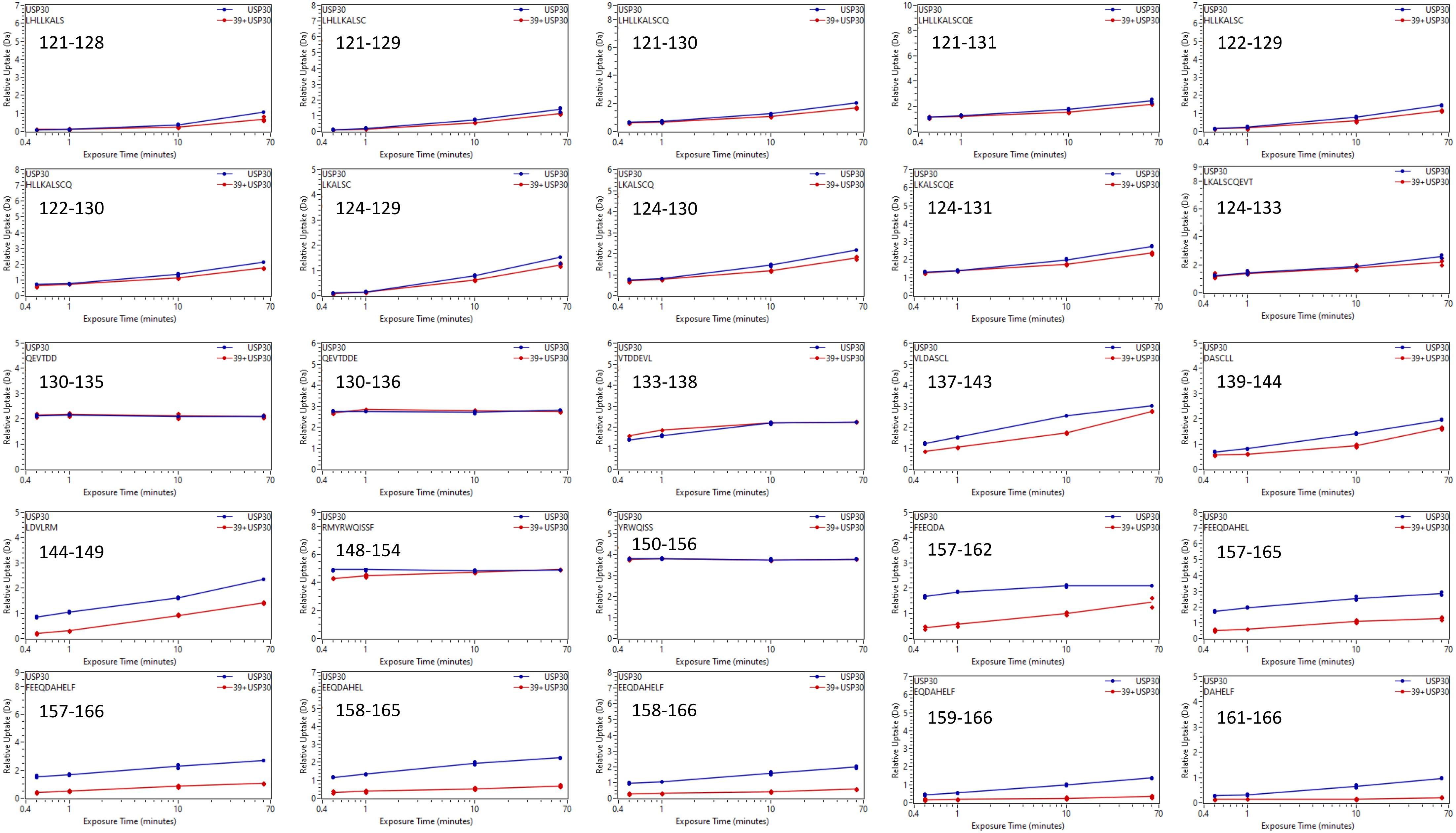

### Supplementary Figure 6 (continued)

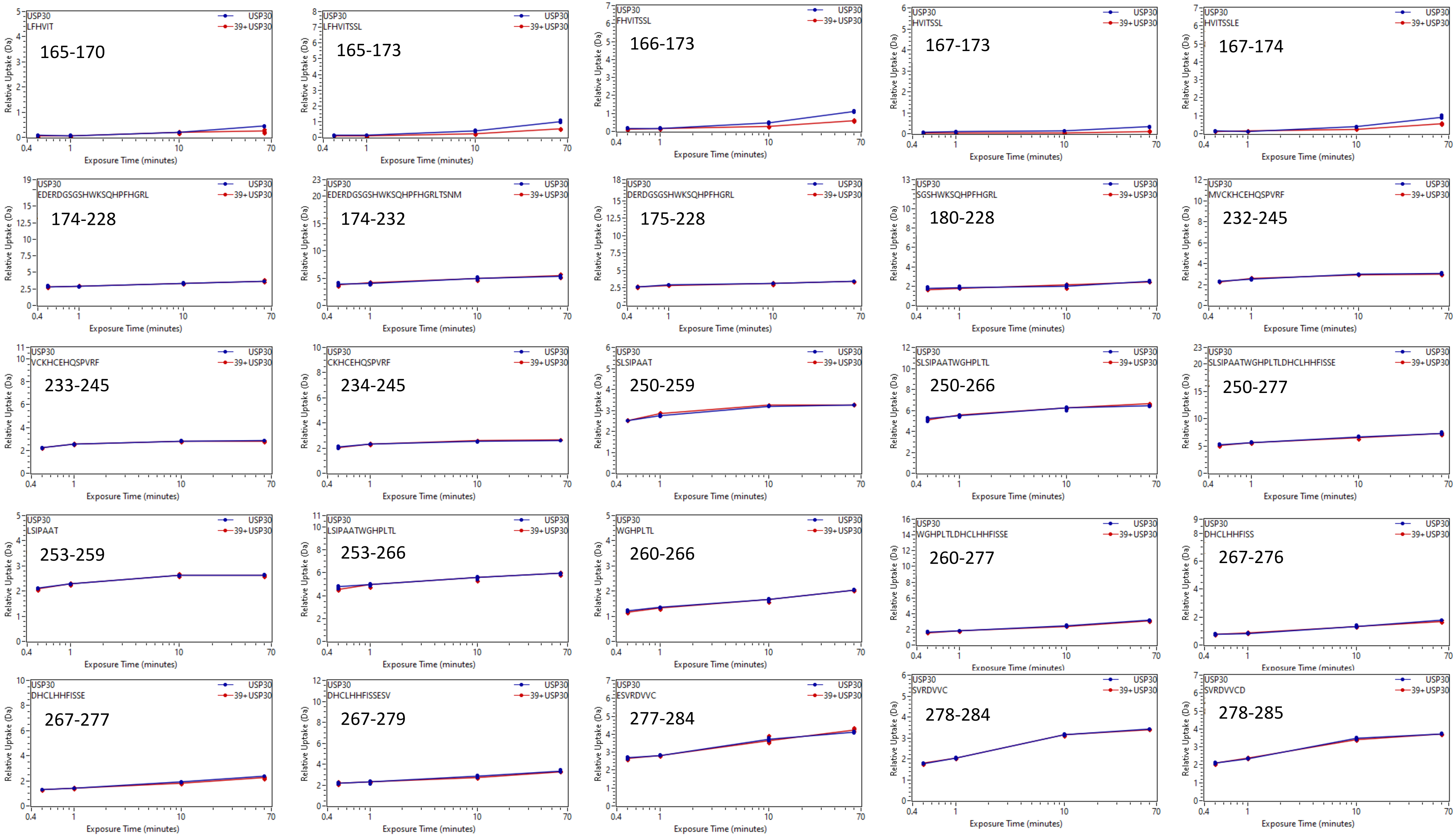

### Supplementary Figure 6 (continued)

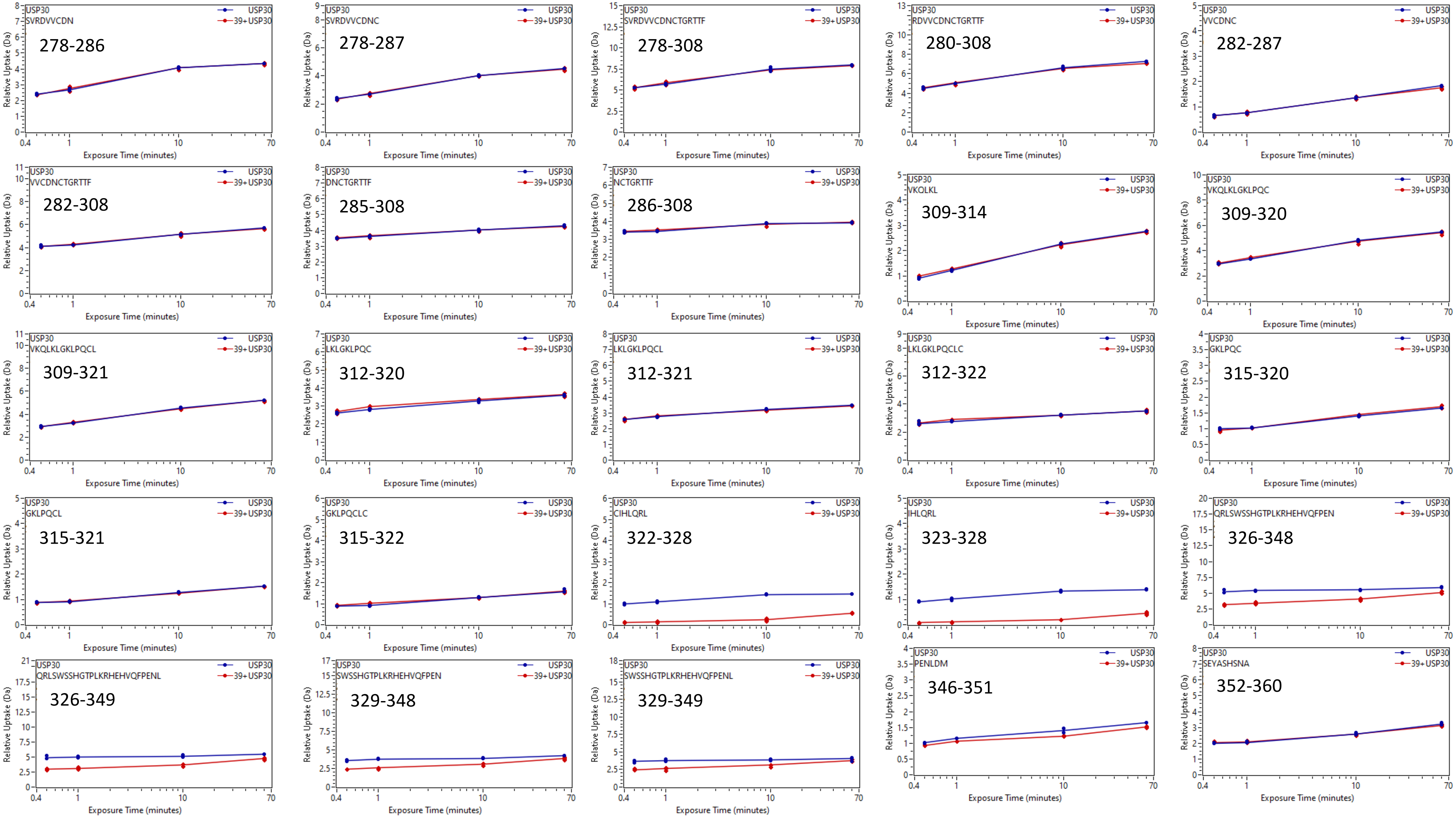

### Supplementary Figure 6 (continued)

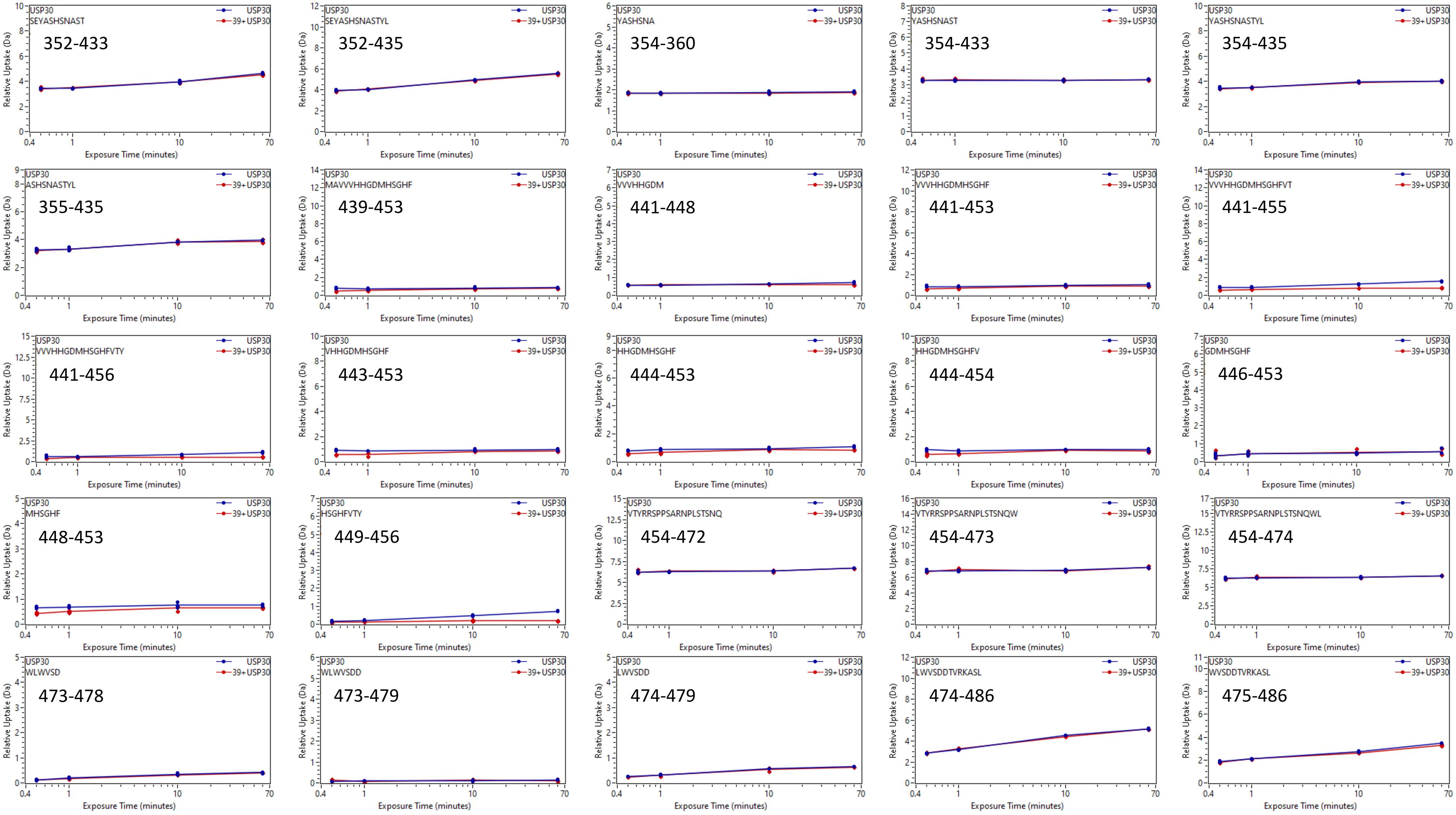

### Supplementary Figure 6 (continued)

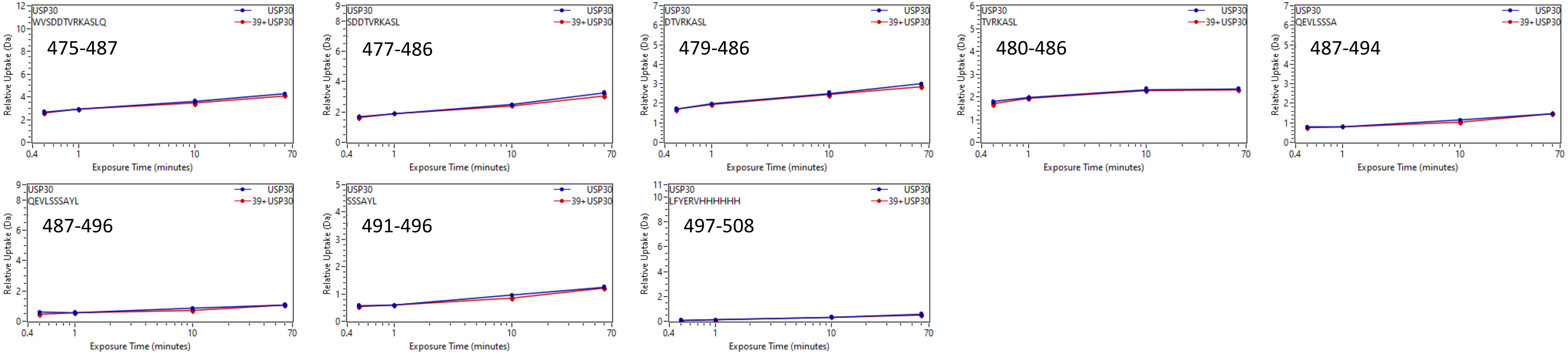
