## Supporting Information for "Structural Premise of Selective Deubiquitinase USP30 Inhibition by Small-Molecule Benzosulfonamides"

Contents:

1. Supporting Text
2. Supporting Figure Legends

### 1. SUPPORTING TEXT

*Kinetic assays – steps to determine slow binding inhibitor constants*

In case of a covalent inhibitor, $IC_{50}$ values can be plotted against incubation time and the resulting curve can be fitted to **Eq.1** to obtain the inhibition constant $K_i$, and the rate of enzyme inactivation $k_{inact}$.

$$\boldsymbol{IC_{50}(t) = K_i\left(1+\frac{[s]}{K_m}\right)\cdot\left(\frac{2-2e^{-\eta IC_{50}\cdot k_{inact}\cdot t}}{\eta IC_{50}\cdot k_{inact}\cdot t}-1\right)}$$

$$\text{Where: } \eta \boldsymbol{IC_{50}} = \frac{\boldsymbol{IC_{50}\ (t)}}{\boldsymbol{K_i\left(1+\frac{[S]}{K_m}\right)+IC_{50}(t)}}$$

**Eq.1**

In order to determine the binding kinetics of a non-covalent slow-tight binding compound progress curves were fitted to Eq.2, where [P] is product formed, $t$ is time, $v_s$ is the steady-state rate of the reaction, $v_i$ is the initial rate of the reaction and $k_{obs}$ is the rate at which the reaction converts from $v_i$ to $v_s$.

$$\boldsymbol{[P] = v_s t + \frac{v_i - v_s}{k_{obs}}[1-\exp(-k_{obs}t)]}$$

**Eq.2**

The resulting $k_{obs}$ values were plotted against concentration and fitted to Eq.3a to obtain $k_5$, $k_6$ and $K_i^{app}$ .

$$\boldsymbol{k_{obs} = k_6 + \frac{k_5[I]}{K_i^{app}+[I]}}$$

**Eq.3a**

Where $k_{obs}$ is the rate at which the reaction converts from $v_i$ to $v_s$, $I$ is concentration of inhibitor, $k_5$ is the forward rate of E-I converting to E-I*, $k_6$ is the reverse rate of E-I converting to E-I* and $K_i^{app}$ is the apparent equilibrium constant for the formation of E-I.

Next, an equation for converting $K_i^{app}$to $K_i$ is used, assuming competitive inhibition (Eq.3b).

$$K_i = \frac{K_I^{app}}{1+(\frac{[S]}{K_m})}$$

**Eq.3b**

Where $K_i^{app}$ is the apparent equilibrium constant for the formation of EI, $K_i$ is the equilibrium constant for the formation of E-I, S is the initial substrate concentration in the reaction and $K_m$ is the Michaelis-Menten constant for USP30 Ub-Rho110.

The value for $K_i^*$ was determined (Eq.3c), which is the equilibrium constant for the formation of E-I*.

$$K_i^* = \frac{K_i k_6}{k_5 + k_6}$$

**Eq.3c**

$K_i$ is the equilibrium constant for the formation of E-I, $k_5$ is the forward rate of EI converting to EI* and $k_6$ is the reverse rate of EI converting to E-I*.

### 2. SUPPLEMENTARY FIGURE LEGENDS:

**Figure S1: Confirmation of USP30 complex formation with Inhibitor 39**. The *m/z* of apo-USP30 and holo-30 (1:1 ratio of protein:compound) was measured over 75 min by Rapidfire MS. No change in *m/z* measurements indicated that the complex was stable and maintained over the time course of our experiment.

**Figure S2: Peptide map of USP30**. Overall, 133 peptic USP30 peptides were selected for differential HDX-MS analysis following digestion of the unlabelled protein with pepsin. This corresponded to a sequence coverage of 96.2%, with an average peptide redundancy of 4.19. Individual peptides are represented by a blue bar on the plot.

**Figure S3: Differential HDX-MS of USP30 and 39**. a) Domain organisation of USP30. b) Relative fractional uptake of apo-USP30 (top) and holo-USP30 (bottom) by HDX-MS. All time points are individually coloured, with each measured USP30 peptide displayed on the x-axis. Peptides are sorted from the USP30 N-terminus to the USP30 C-terminus. Deuterium incorporation is mapped to the y-axis. c) Differential deuterium uptake between holo- and apo-USP30. Regions of greatest perturbation are labeled I-IX.

**Figure S4: Architecture of USP30 inferred by HDX-MS.** The HDX-MS data was mapped to the crystal structure (5OHK) of both apo- and holo-USP30. Dynamic HDX behavior indicative of structural elements are colored green, while their disordered counterparts are colored grey. Regions are blocked and completely solvent inaccessible are coloured purple.

**Figure S5: Temporal exchange of USP30**. The HDX-MS data was mapped to USP30 at each time point and the magnitude of perturbation induced by inhibitor 39 binding to USP30 color-coded. The majority of the protein is unaffected by the protein, but several regions undergo substantial (>27.5%) solvent shielding in the presence of the compound.

**Figure S6: HDX-MS uptake curves for individual USP30 peptides.** Uptake curves for apo-USP30 are colored blue, while those of holo-USP30 are colored red.
